# Convolutional architectures are cortex-aligned de novo

**DOI:** 10.1101/2024.05.10.593623

**Authors:** Atlas Kazemian, Eric Elmoznino, Michael F. Bonner

**Author notes:** **Please address correspondence to:** Atlas Kazemian or Michael F. Bonner Department of Cognitive Science Johns Hopkins University 3400 N. Charles Street Baltimore, MD 21218.

## Abstract

What underlies the emergence of cortex-aligned representations in deep neural network models of vision? Earlier work suggested that shared architectural constraints were a major factor, but the success of widely varied architectures after pre-training raises critical questions about the importance of architectural constraints. Here we show that in wide networks with minimal training, architectural inductive biases have a prominent role. We examined networks with varied architectures but no pre-training and quantified their ability to predict image representations in the visual cortices of monkeys and humans. We found that cortex-aligned representations emerge in convolutional architectures that combine two key manipulations of dimensionality: compression in the spatial domain, through pooling, and expansion in the feature domain by increasing the number of channels. We further show that the inductive biases of convolutional architectures are critical for obtaining performance gains from feature expansion—dimensionality manipulations were relatively ineffective in other architectures and in convolutional models with targeted lesions. Our findings suggest that the architectural constraints of convolutional networks are sufficiently close to the constraints of biological vision to allow many aspects of cortical visual representation to emerge even before synaptic connections have been tuned through experience.

## INTRODUCTION

Understanding the computations that transform patterns of light on the retina into representations that lead to complex behavior is a longstanding challenge in neuroscience. Early work focused on theory-guided, hand-engineered models for simulating image-evoked neural responses, but these models had limited success beyond primary visual cortex (Carandini et al., 2005; Jones & Palmer, 1987; Movshon et al., 1978; Riesenhuber & Poggio, 1999). Following the advent of deep learning, it was discovered that deep neural networks (DNNs) could achieve unprecedented performance in predicting image-evoked responses in the ventral visual stream, and they have since become the leading models of visual computation in the brain (Agrawal et al., 2014; Khaligh-Razavi & Kriegeskorte, 2014; Yamins et al., 2014). However, despite over a decade of research, it remains an open question what key factors underlie the representational similarities between DNNs and visual cortex (Chen & Bonner, 2025; Conwell et al., 2022; Elmoznino & Bonner, 2024; A. Saxe et al., 2020; Serre, 2019; Storrs et al., 2021).

A prominent theory proposes that cortex-aligned representations emerge in DNNs when their constraints and objectives match those of biological vision (Cao & Yamins, 2021a, 2021b). This theory builds on foundational work in neuroscience arguing that biological visual representations are strongly constrained by neuroanatomic architecture and optimized for specific ethologically relevant objectives (R. Cao & Yamins, 2021b; Konkle & Alvarez, 2022; Kriegeskorte, 2015; Richards et al., 2019; Yamins & DiCarlo, 2016). However, recent findings have challenged this constraint-based perspective. Most importantly, it has been shown that DNNs with widely varied architectures and training objectives predict ventral-stream representations equally well after being pre-trained on massive datasets (Conwell et al., 2022; Storrs et al., 2021). This suggests that for highly trained DNNs, architectural and objective-based constraints are relatively weak (Chen & Bonner, 2025; Conwell et al., 2022; Elmoznino & Bonner, 2024). There is, thus, a crucial and unresolved question: How can we reconcile the apparent degeneracy of DNN designs with the strong role that architectures and task objectives are thought to play in shaping the representations of biological vision?

In this article, we present a different perspective on this problem by examining how the encoding performance of untrained DNNs changes as a function of targeted architectural manipulations. By removing the influence of massive pre-training, our approach reveals the surprising extent to which architectural factors alone can promote the emergence of cortex-aligned representations. Specifically, we scaled the number of random features in convolutional, fully connected, and transformer architectures and quantified their ability to predict image-evoked responses in the visual cortices of monkeys and humans. Our results showed a strong effect of scaling that was specific to convolutional architectures. Scaling the deeper layers of a convolutional network yielded striking performance gains—the best model even approached the performance of a classic pre-trained network in the monkey data. In contrast, scaling the features of fully connected and transformer architectures yielded only small improvements, resulting in substantially worse performance than the convolutional architecture despite being matched on dimensionality. Further, the performance of the convolutional architecture depended critically on spatial locality and nonlinear activation functions, again demonstrating that the benefits of scaling were highly architecture-dependent.

Together, these findings show that the architectural biases imbued into convolutional networks allow many aspects of cortical visual representation to readily emerge even before synaptic connections have been tuned through experience. Thus, while pre-training on large-scale datasets may be sufficient to induce cortex-aligned representations in many degenerate architectures, the representations of convolutional architectures exhibit a remarkable degree of cortical alignment in their initial random states.

## RESULTS

### Scaling untrained neural networks to predict visual cortex representations

We examined how the encoding performance of untrained neural networks changed when scaling the dimensionality of their features. We focused on convolutional, fully connected, and transformer architectures. The convolutional architecture, which we refer to as the Expansion model, contained a hierarchy of layers that each perform the following operations: convolution, nonlinear activation, and pooling (Figure 1a). The first layer contained a pre-defined set of wavelet filters (Bruna & Mallat, 2012; Pogoncheff et al., 2023; Yue et al., 2014, 2020) but all other layers contained randomly initialized weights. Our key manipulation was the number of random convolutional filters in the last layer. We also analyzed fully connected and transformer architectures and manipulated the number of randomly initialized features in their last layers. Detailed descriptions of network architectures can be found in the Methods.

**Figure 1.**
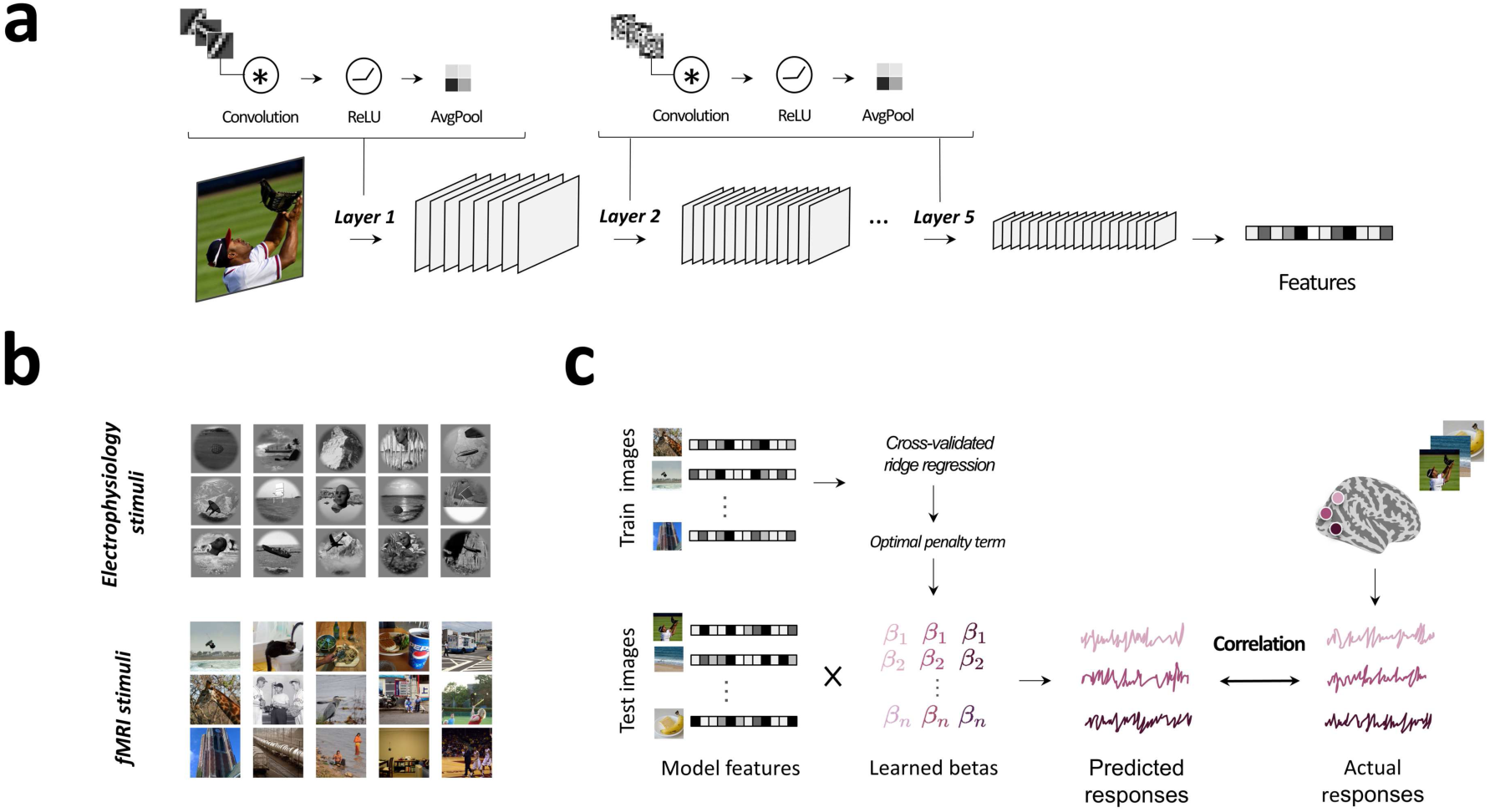
Convolutional model architecture and evaluation framework. **a)** Our convolutional architecture consists of a repeating set of modules that are stacked in a hierarchy of five layers. In the first layer, images are convolved with a set of pre-defined wavelet filters, then passed through a rectified linear unit (ReLU) activation function, and an average pooling operation. In the deeper layers, inputs are convolved with a large set of randomly initialized filters, followed by ReLU and average pooling. Features were extracted from the last layer of the network and used to evaluate cortical encoding performance. **b)** The top panel shows examples of stimuli from the monkey electrophysiology dataset (Majaj et al., 2015), and the bottom panel shows examples of stimuli from the human fMRI dataset (Allen et al., 2022). **c)** To evaluate cortical encoding performance, model image representations were linearly mapped to image-evoked cortical responses using ridge regression. Regression coefficients and the optimal ridge penalty parameter were estimated on a set of training images and performance was evaluated on held-out test images. Encoding performance scores were obtained by computing correlations between the predicted and actual cortical responses in the test set.

We quantified the ability of these networks to predict image-evoked responses in visual cortex in both monkey electrophysiology and human fMRI data. The monkey dataset consists of single-unit recordings from two macaques who viewed 3200 images of objects on synthetic backgrounds (Majaj et al., 2015). We considered all units in V4 and IT regions from both macaques. The second dataset, called The Natural Scenes Dataset (NSD) (Allen et al., 2022), is the largest existing human fMRI dataset on natural scene perception, containing responses to ∼73,000 images across 8 participants. We considered fMRI responses from three regions of interest (ROIs) from all participants: early, midventral, and ventral visual streams. Example stimuli from both datasets are shown in Figure 1b. Using regularized linear regression, we fit encoding models that mapped the final layer of each neural network to the responses of electrophysiology units or fMRI voxels. These encoding models were estimated on a set of training images, and their performance was evaluated on a held-out set of test images. Encoding performance was measured as the correlation between predicted and actual cortical responses in the test set (Figure 1c). Additionally, Extended Data Figure 1 shows a replication of our key findings in a third dataset (the large-scale THINGS fMRI dataset, which contains responses to thousands of object images) (Hebart et al., 2023).

We examined encoding performance as a function of dimensionality expansion in the output units of neural networks. Specifically, we sought to i) determine if dimensionality expansion could substantially improve encoding performance, ii) determine how much of a performance gain might be obtained, and iii) determine whether these potential gains differed across architectures. Our goal was to characterize the point at which encoding performance converged and no longer improved from the addition of more dimensions. We, therefore, increased the number of random features in the last layer by multiple orders of magnitude and quantified encoding performance at each level of dimensionality.

In all ROIs in both the monkey and human data, the results show two main trends (Fig. 2). First, the convolutional architecture outperformed the other architectures across all levels of dimensionality. Second, all three architectures obtained performance gains from dimensionality expansion, but interestingly, the degree of the performance gain was markedly different for the convolutional architecture compared with the other two architectures, despite being matched on dimensionality. Specifically, we found that dimensionality expansion strongly improved the performance of the convolutional architecture while resulting in substantially smaller improvements for the fully connected and transformer architectures. Together, these findings indicate that the inductive biases of the convolutional architecture strongly promote the emergence of cortex-aligned representational dimensions, which can be readily discovered through simple linear transformations of a random convolutional feature space. Note, however, that these findings do not suggest that visual cortex itself is a wide random feature bank. As shown through representational similarity analysis in Extended Data Figure 2, the use of a linear transformation is crucial for identifying the subset of dimensions in these wide networks that best align with visual cortex.

**Figure 2.**
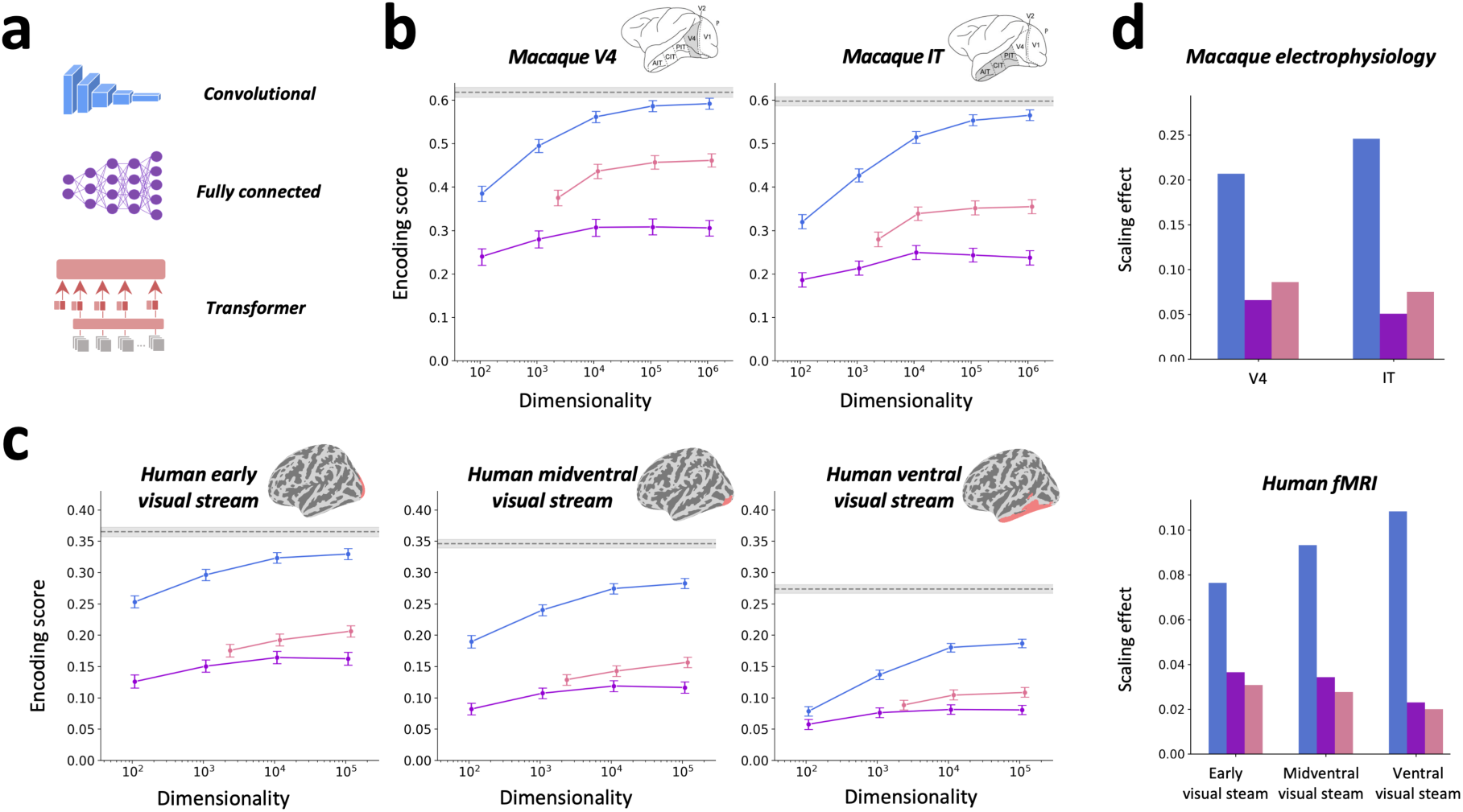
Dimensionality expansion strongly improves the encoding performance of convolutional neural networks. **a)** The effects of dimensionality expansion were examined for a convolutional neural network, a deep fully connected network, and a vision transformer. **b)** These plots illustrate the encoding performance for all three architectures as a function of dimensionality expansion. There was no pre-training for these networks. Encoding performance was evaluated for regions V4 and IT in the monkey electrophysiology data. The x-axis plots the number of random features in the output layer, and the y-axis shows the encoding score for predicting image-evoked cortical responses. The gray dashed line indicates the performance of the best performing convolutional layer of pre-trained AlexNet. The convolutional architecture without pre-training attained large performance gains as a function of dimensionality expansion and approached the performance of pre-trained AlexNet. In contrast, the other two architectures showed much less improvement as the number of output features was expanded. The encoding performance plots in this panel and in panel c show the mean performance across units/voxels from all participants, and the error bars denote 97.5% confidence intervals from 1,000 bootstrap samples. **c)** These plots show the same analyses as in panel b but for regions along the ventral stream in the human fMRI data. The encoding performance of the largest convolutional model approaches that of pre-trained AlexNet in the early visual stream and reaches ∼70% of the performance of pre-trained AlexNet in the higher-level ventral stream. **d)** A summary statistic for the effect of dimensionality expansion was calculated as the difference in performance between the highest- and lowest-dimensional version of each architecture. The performance of the convolutional model was strongly modulated by the dimensionality of its random feature space. In contrast, the effect of dimensionality expansions was much weaker in the other two architectures.

For comparison, we also plotted the performance of a conventional CNN architecture (AlexNet) that was pre-trained on ImageNet classification. We found that in both the V4 and IT regions of monkey visual cortex, the largest untrained convolutional architecture approached the performance of pre-trained AlexNet, demonstrating the remarkable degree to which high-level object representations in monkey visual cortex can be predicted from random convolutional features without goal-driven pre-training (Figure 2b). Interestingly, in the human data, we observed a larger performance gap between the untrained convolutional architecture and AlexNet, and we found that this performance gap increased along the hierarchy of the ventral stream (Figure 2c).

Extended Data Figure 3 shows additional comparison models, including the current state-of-the-art model for the monkey IT data on the BrainScore platform, BarlowTwins (Schrimpf et al., 2018). The BarlowTwins model obtains a higher encoding score than AlexNet in monkey IT (mean r-value = 0.67 for BarlowTwins vs. 0.60 for AlexNet) but is comparable to AlexNet in the human ventral stream (mean r-value = 0.29 for BarlowTwins vs. 0.27 for AlexNet). As shown in Extended Data Figure 3, none of the models that we tested reached an estimate of an upper bound for the noise ceiling in any ROI in both the monkey data (mean noise ceiling r-value = 0.88 for V4 and 0.86 for IT) and human data (mean noise ceiling r-value = 0.47 in the early visual stream, 0.48 in the midventral visual stream, and 0.37 in the ventral visual stream).

In follow-up analyses using variance partitioning, we found that the explained variance of the Expansion model is shared with AlexNet (Extended Data Figure 4). Thus, the Expansion model obtains high encoding scores by accounting for variance that is shared with a pre-trained supervised network, as opposed to explaining variance that is unique to untrained networks.

Finally, we performed an analysis to examine how the expansion of random networks affected the latent dimensionality of their image representations. We computed a metric of effective dimensionality by counting the number of principal components that accounted for 85% of the variance in each network’s image representations, and we then plotted encoding performance as a function of this dimensionality metric. As shown in Extended Data Figure 5, the results reveal several key trends. First, in all architectures, we see that effective dimensionality generally increases as the number of features is increased, but in the monkey data it reaches an upper bound, after which further expansions of the network have a negligible impact on effective dimensionality. Second, as noted in previous work (Elmoznino & Bonner, 2024), we see that effective dimensionality alone is not sufficient to explain encoding performance, as demonstrated by the performance differences observed across different architectures with the same effective dimensionality. Third, and most importantly, we see that the encoding performance of each network increases as a function of its effective dimensionality, suggesting that feature expansion is only useful insofar as it increases the effective dimensionality of a network’s natural image representations. These results raise the intriguing question of whether it may be possible to yield further gains in encoding performance through architectural manipulations that increase effective dimensionality beyond the levels reached here.

### Emergent properties of untrained convolutional neural networks

The large performance gains of the expanded convolutional network suggest that brain-relevant representations readily emerge when natural images are processed with random convolutional filters. We further explored the emergent properties of untrained CNNs in two follow-up analyses. First, we performed principal component analysis (PCA) on the image representations of the CNNs to identify their principal modes of variance for natural images, and we found that almost all the explained variance of the high-dimensional CNNs could be accounted for by a reduced set of principal components (PCs) (Figure 3). These analyses also show that there is a benefit from first performing dimensionality expansion using a large set of random convolutional filters before reducing the dimensionality through PCA. For example, when the model containing 10^6^ random features was reduced to 10^2^ PCs, its performance was still substantially better than a model with 10^2^ random features, even though both models have the same number of regressors. This is because some of the features in these random networks will be redundant with others and some will capture little-to-no variance across natural images. This means that the number of useful dimensions in a network is not equal to the number of neurons it contains (Casper et al., 2021). As a result, the dimensionality of a wide network can be reduced through PCA with little impact on its encoding performance, and by the same logic, a narrow network may perform worse than a PCA-reduced network with the same number of dimensions.

**Figure 3.**
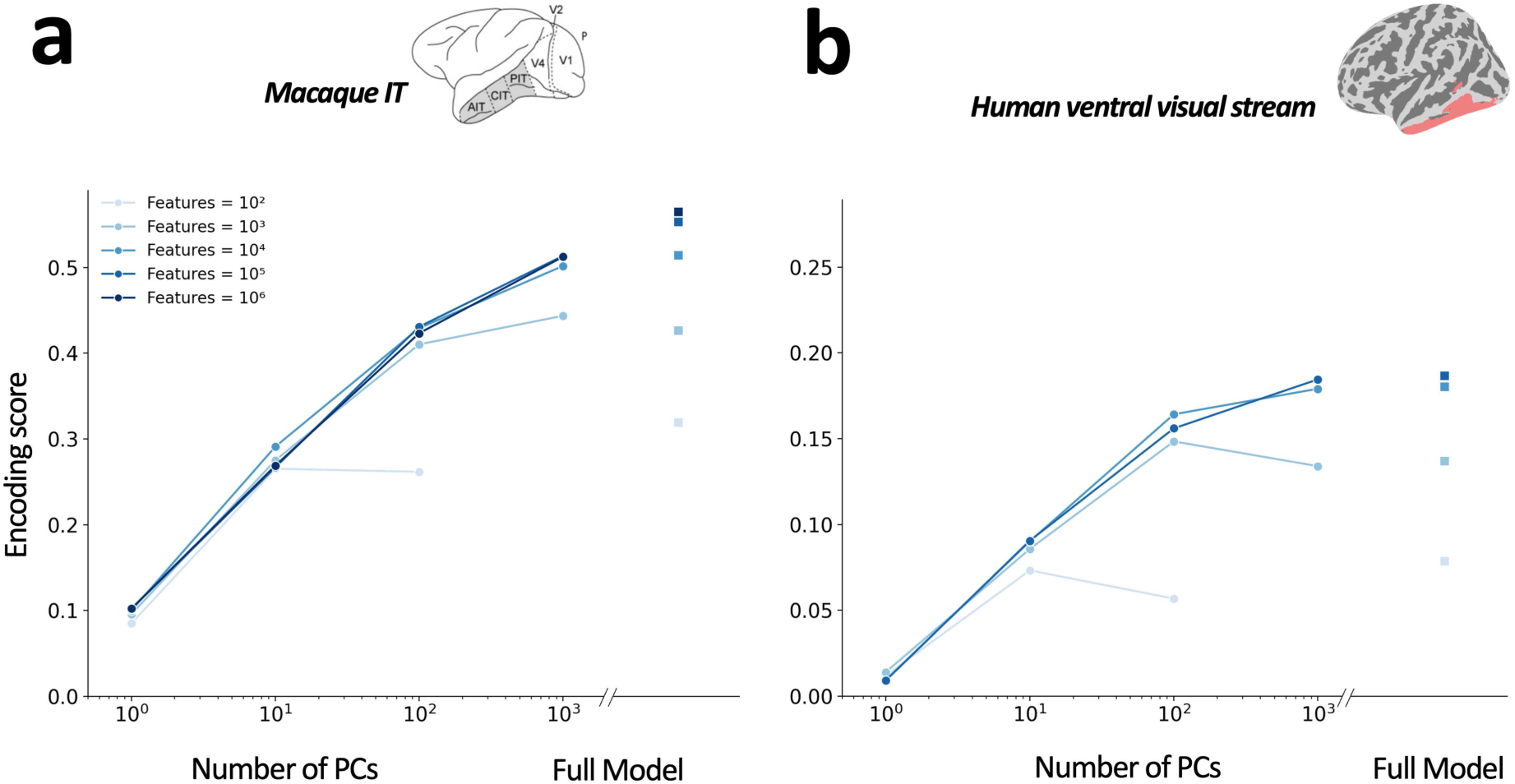
Encoding performance remains high after dimensionality reduction. These plots show encoding performance for models whose dimensionality was reduced through principal component analysis. Networks with high-dimensional random feature spaces in their last layer were reduced to a smaller set of principal components (PCs). PCs were identified using an independent set of stimuli, and they reflect the principal directions of variance for the natural image representations of each network. Results are shown for the monkey IT region and the human high-level ventral stream. Panels **a** and **b** show encoding performance for the convolutional networks from Figure 2 after their final layers were reduced to varying numbers of PCs. An inset on the right of each plot shows the performance of the original networks without dimensionality reduction. Color shades represent the number of random features in the last convolutional layer of each network. Both plots show that all networks can be reduced to a lower-dimensional subset of PCs with only a minor drop in encoding performance. For example, the dimensionality of the largest networks can be reduced by orders of magnitude but retain nearly the same level of encoding performance.

We next performed a visualization analysis to explore the representational space of our largest untrained CNN. Specifically, we visualized the image representations of the CNN in a 2D space using t-distributed stochastic neighbor embedding (tSNE). This visualization shows that despite having no pre-training for classification, the CNN forms many intuitively interpretable clusters of images, including clusters related to sports, vehicles, food, animals, and people (Figure 4). We confirmed that these embeddings contain a significant degree of semantic clustering by computing the average within-category versus between-category distances using the object category labels for all images. As expected, this categorical clustering index was significantly above chance (p<0.001, permutation test). This suggests that there is a remarkable degree to which images naturally organize into semantically meaningful clusters in the representational space of high-dimensional untrained CNNs.

**Figure 4:**
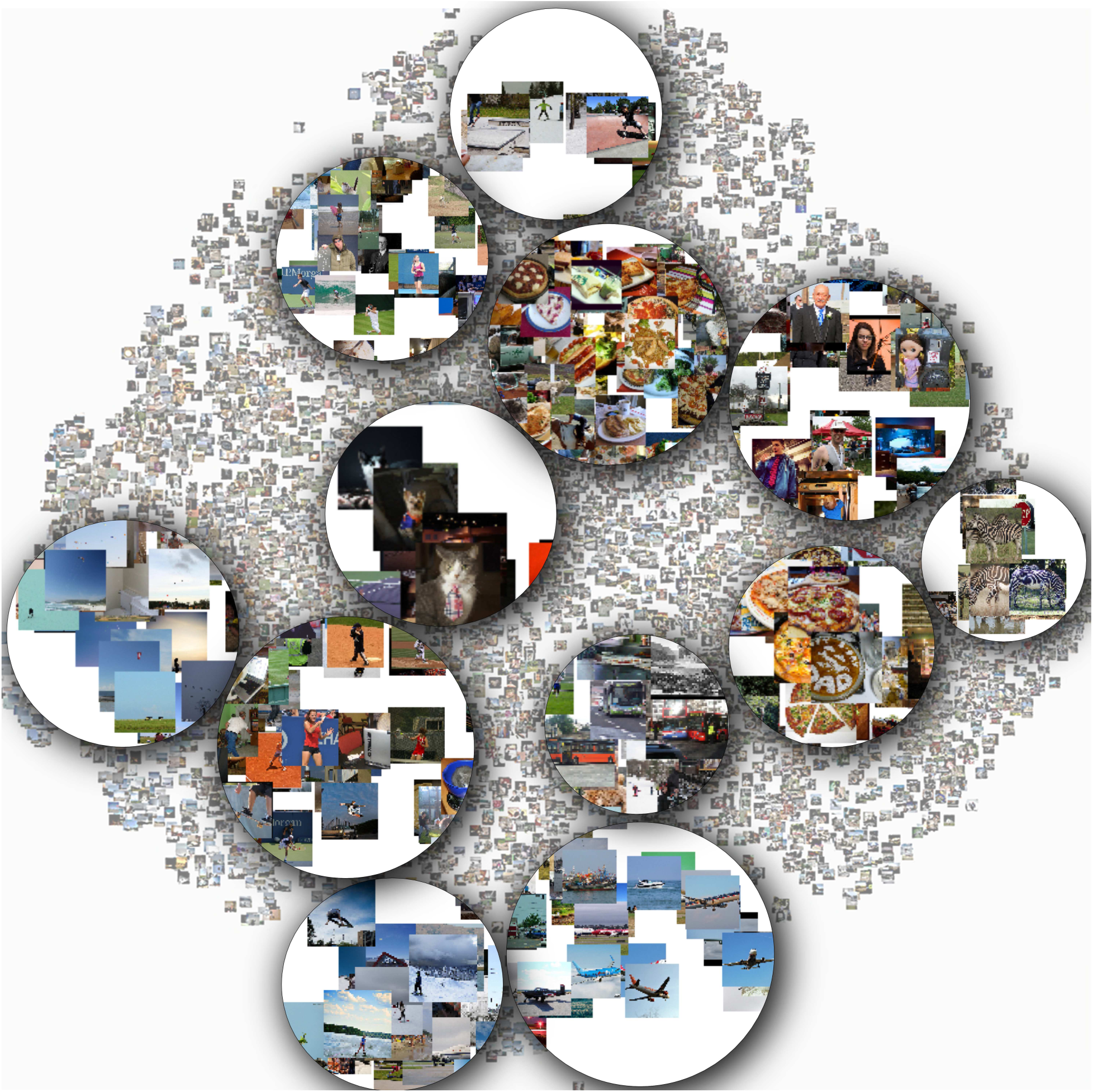
Two-dimensional visualization of image representations in a high-dimensional untrained convolutional network. Image representations from the final layer of a high-dimensional untrained convolutional network were embedded in two dimensions using t-distributed stochastic neighbor embedding. These embeddings show that even in an untrained network, images naturally form clusters that are intuitively interpretable. This visualization highlights several of these clusters, including clusters related to sports, vehicles, food, animals, and people.

### Critical architectural components

Our analyses thus far show that strong encoding performance cannot be obtained through dimensionality expansion in arbitrary architectures but, instead, depends critically on the inductive biases of the convolutional architecture. We next sought to determine which components of the convolutional architecture are most important for its performance. We performed a series of analyses in which we altered specific properties of the architecture and evaluated encoding performance in high-level visual cortex.

We first confirmed that nonlinearities were critical by removing the nonlinear activation function in all layers while keeping the rest of the architecture the same (Figure 5). As expected, we found that in both the monkey and human data, performance was worse in a network without nonlinearities and there was little-to-no benefit of dimensionality expansion. This result demonstrates that a nonlinear transformation is needed to explain the representations of high-level visual cortex.

**Figure 5:**
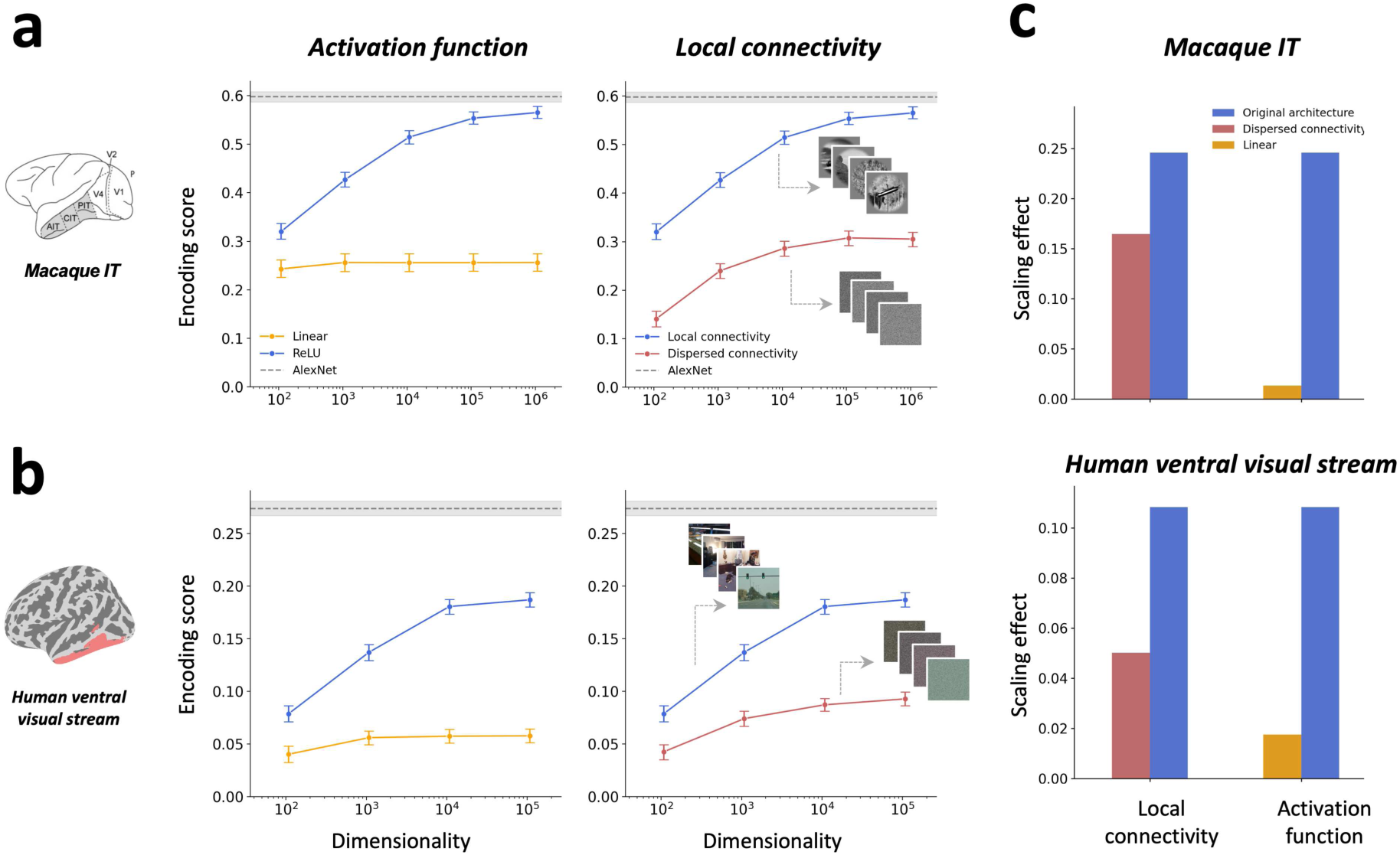
Analysis of critical architectural components of the convolutional network. **a-b)** These plots show the performance of the largest untrained convolutional network after removing key architectural components but keeping all other architectural properties intact. Panel a shows results for macaque IT, and panel b shows results for the human high-level ventral stream. The left plots show that removing nonlinearities results in a large drop in encoding performance in both the monkey and human data, and it prevents the model from improving as a function of dimensionality expansion. The right plot shows that removing the spatial locality of the convolutional filters through fixed permutations of image pixels results in an overall decrease in encoding performance, though networks with such dispersed connectivity still have moderate benefits from dimensionality expansion. Example images are shown to illustrate the effect of applying fixed permutations to image pixels. Plotting conventions are the same as in Figure 2. **c)** These plots show the effects of dimensionality expansion, as in Figure 2d. They demonstrate that without nonlinearities, the benefits of dimensionality expansion are removed and without spatial locality, they are diminished.

We next examined whether the spatial locality of the convolutional filters was important for model performance (Figure 5). Spatial locality is one of the central inductive biases of convolutional neural networks, and it is known to reduce the curse of dimensionality for deep learning of image representations (Poggio et al., 2017). Thus, we expected it to have a critical role in encoding performance. To examine this question, we applied a fixed permutation to image pixels by shuffling the pixel locations in the same way for each image. This made the inputs to the convolutional filters spatially distributed rather than spatially local (i.e., each convolutional filter operated on a collection of pixels from across the image rather than from a local image patch). All other architectural properties remained unchanged. As expected, we found that in both the monkey and human data, there was a large drop in encoding performance in the network without spatially local image representations, demonstrating that spatial locality is critical for achieving strong encoding performance.

We additionally performed a series of follow-up experiments to examine the effects of architectural changes applied to only the final layer of the network instead of all layers (Extended Data Figure 6). The findings show that even in the highest layer of the network it is beneficial to compute spatially local representations. They also show that ablating the nonlinearity in the final layer has little effect, which means that the crucial nonlinear operations are those that occur in earlier layers.

Further analyses of other architectural factors are reported in Extended Data Figures 7-9. Briefly, these analyses show that there is a small benefit from the use of pre-defined wavelet filters in the first layer. For example, in the monkey IT data the largest model reached a mean r-value of 0.57 with pre-defined wavelets vs. 0.50 with random filters, and in the human ventral stream the largest model reached a mean r-value of 0.19 with pre-defined wavelets vs. 0.15 with random filters. These analyses also show that multiple types of nonlinear activation functions and random initialization methods yield similar patterns of results.

In sum, the findings from these architectural manipulations highlight the effectiveness of the convolutional network’s central inductive biases. Specifically, without nonlinear transformations and spatially local image representations, dimensionality expansion is insufficient to achieve strong encoding performance.

### Encoding performance can be strong even when classification accuracy is weak

Pre-training neural networks on image classification improves their cortical encoding performance, and it has been argued that strong classification performance is a signature of high-performing neural network models of visual cortex (R. Cao & Yamins, 2021b; Khaligh-Razavi & Kriegeskorte, 2014; Yamins & DiCarlo, 2016). However, our findings show that even untrained convolutional networks can achieve good encoding performance when they are high-dimensional and have the right architectural properties, suggesting that classification performance may not be as tightly linked to encoding performance as previous work suggests. To address this question, we examined the classification performance of our largest untrained convolutional model, which we call the Expansion model here, and we compared it with AlexNet pre-trained on ImageNet. We used the Places365-Standard dataset, which contains images from 365 different scene categories (Zhou et al., 2018) (Figure 6a). We used Places rather than ImageNet so that we could evaluate both models on a dataset on which they had not previously been optimized. We fit linear classifiers to the final convolutional layer of each network, and we evaluated performance on held-out images. Figure 6b shows that the Expansion model performs much worse than AlexNet at classifying scene categories. Thus, even though the encoding performance of the Expansion model is strong enough to rival AlexNet in monkey visual cortex, it is, nonetheless, much worse than AlexNet at classifying images. This suggests that optimization for image classification is not critical for obtaining highly predictive encoding models of visual cortex.

**Figure 6:**
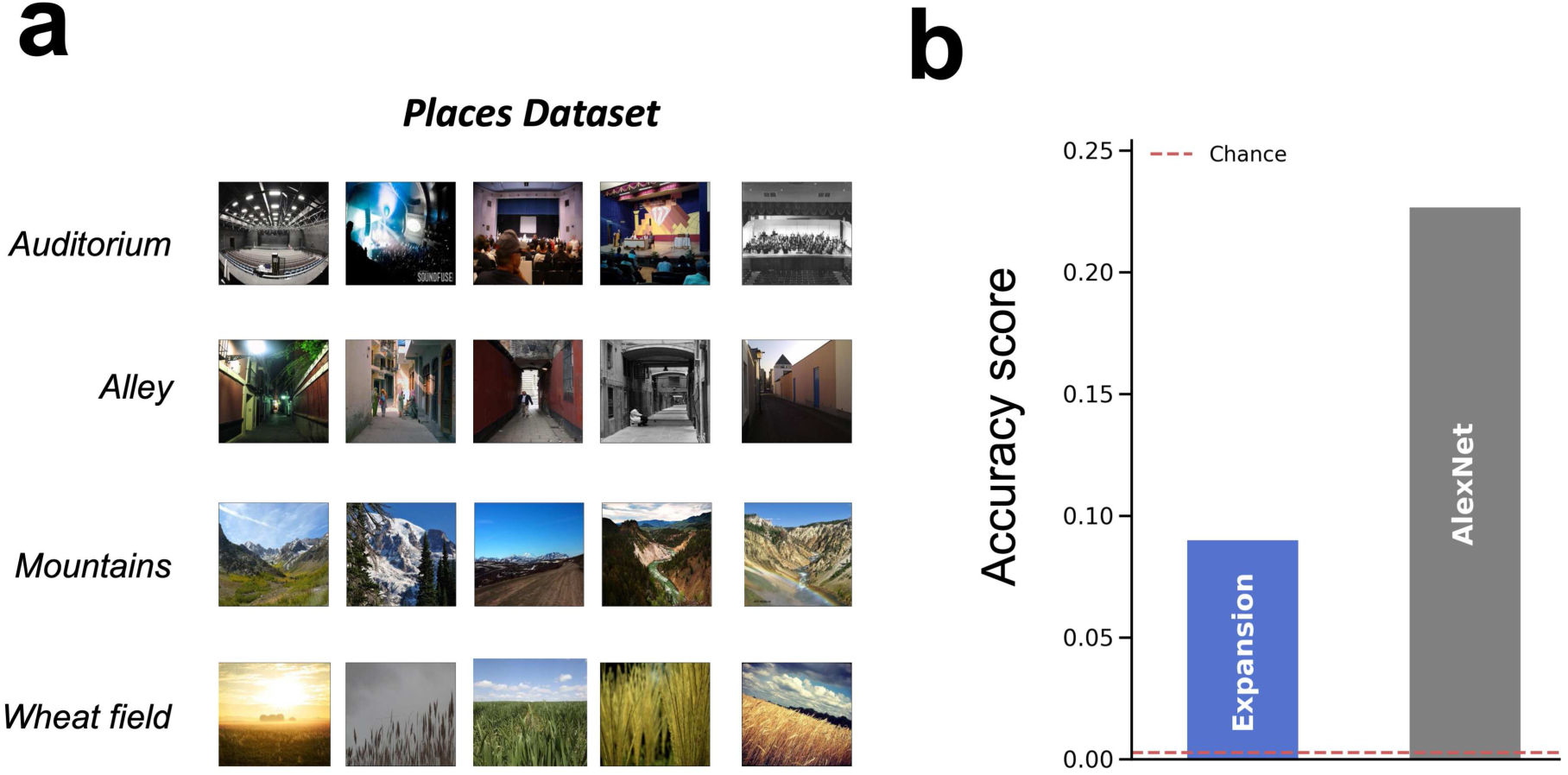
Image classification performance for the untrained convolutional network. **a)** Performance was evaluated for classification on the 365 scene categories in the Places dataset (Zhou et al., 2018). **b)** A 365-way linear classifier was fit to the principal components of features from the final convolutional layer of the Expansion model with 10^5^ features, and AlexNet pre-trained on ImageNet. Classification accuracy was evaluated on held-out test images. This plot shows that the Expansion model performs poorly at image classification despite obtaining strong encoding performance scores in monkey visual cortex and early to mid-level regions of human visual cortex.

## DISCUSSION

We show that dimensionality expansion in an untrained convolutional neural network achieves surprisingly strong performance at explaining image-evoked responses in the primate visual cortex, in some cases reaching the performance of a standard pre-trained network. This performance is not simply an effect of model scale, where any high-dimensional model would successfully predict brain responses. Instead, it reflects the effectiveness of the convolutional architecture’s inductive biases, which greatly increase the likelihood of discovering brain-relevant representational dimensions through random feature sampling.

Previous work has argued that task-based pre-training is among the most important factors for obtaining brain-aligned representations in DNNs (R. Cao & Yamins, 2021b; Conwell et al., 2022; Khaligh-Razavi & Kriegeskorte, 2014; Storrs et al., 2021; Yamins et al., 2014; Yamins & DiCarlo, 2016). Although one of the seminal papers from this line of work examined a variety of untrained network architectures (Yamins et al., 2014), they found that architectural manipulations alone were insufficient to yield strong encoding scores, and they argued that IT-like representations emerge when the right class of architecture is combined with task optimization. In contrast, our findings show that with the right inductive biases, architectural manipulations alone can yield large improvements in encoding performance without pre-training or backpropagation. In fact, in the monkey visual cortex data, these architectural manipulations were sufficient to reach the performance of AlexNet pre-trained on ImageNet. Thus, architectural factors alone can play a powerful role in inducing brain-aligned representations in DNNs.

Even though AlexNet is no longer state-of-the-art, it played a central role in early work that convinced neuroscientists to pursue task-based pre-training as a primary factor in model development (Khaligh-Razavi & Kriegeskorte, 2014; Yamins & DiCarlo, 2016), and it inspired over a decade of computational-neuroscience research that has focused heavily on the study of pre-trained DNNs from the engineering field. This focus on task-based pre-training was a major shift from earlier modeling work, which had a much stronger theoretical emphasis on architectural and computational motifs (Oliva & Torralba, 2001; Portilla & Simoncelli, 2000; Riesenhuber & Poggio, 1999). We wonder if untrained computational models might have received more attention had we known ten years ago that AlexNet-level encoding performance in macaque IT could be reached by an untrained architecture. Looking forward, we suggest that architecture optimization in untrained or minimally trained networks is a promising future direction for exploring the inductive biases that may underlie biological vision.

The performance advantage that we observed for our convolutional architecture compared with other architectures contrasts with recent work showing that pre-trained DNNs with highly distinct architectures can obtain similar levels of encoding performance in visual cortex (Conwell et al., 2022). However, it is important to note that massive pre-training may be sufficient to overcome a lack of brain-aligned inductive biases in diverse network architectures, such as in vision transformers. In fact, previous interpretability studies of vision transformers suggest that they implicitly learn to attend to increasingly large regions across layers, thus resembling the increasing receptive fields of convolutional networks (Cordonnier et al., 2020; Dosovitskiy et al., 2021). Furthermore, even fully connected neural networks trained on translation-invariant data automatically develop convolution-like operations, characterized by local receptive fields and weight sharing (Ingrosso & Goldt, 2022). We suggest that the role of architecture can be better understood by focusing on settings in which the contribution of pre-training has been minimized.

Our results also suggest that previous work does not paint a fair picture of how far we can get in predicting visual cortex responses without gradient descent optimization. This is because many of the previously studied architectures were specifically designed to obtain strong benefits from deep learning. It is, thus, not surprising that when comparing trained and untrained versions of these architectures, there is a major advantage of training. Our work examines architectural variations that would be difficult to train through gradient descent due to the large dimensionality of their feature spaces. In doing so, we show that untrained CNNs can perform far better than prior research indicated.

Previous work has examined the surprising effectiveness of feature representations in randomly initialized convolutional networks. Jarrett et al., 2009 showed that a single-layer, randomly initialized CNN competes with an end-to-end trained network in classifying objects from the Caltech 101 dataset. This finding led Saxe et al., 2011 to investigate why random weights performed so well in such a case. To this end, they showed that a combined convolution and pooling architecture can yield spatial frequency selectivity and translation invariance using only random filters. More recently, Cao & Wu (2021) showed that a random, untrained CNN has an inductive bias to localize objects. They suggest that this phenomenon may be driven by the nature of images, where the background is relatively texture-less compared to the foreground objects, increasing the chance that the background will be deactivated by ReLU.

Studying the potential of untrained networks has also gained traction in the field of neuroscience. In an effort to test the effectiveness of untrained CNNs in predicting neural responses in the visual cortex, Baek et al. (2021) evaluated the performance of untrained AlexNet in explaining fMRI responses to faces and showed that face-selective units arise within the network in the absence of supervised training (Baek et al., 2021). In a different study, to improve the biological plausibility of current encoding models, Gieger et al. (2022) investigated model performance under minimized supervised training. Using a biologically inspired CNN with recurrent connections, they showed that training only around 5% of model synapses results in an 80% match to the performance of the fully trained network at modeling macaque ventral stream (Geiger et al., 2022). Studies of the mouse visual system have also demonstrated the surprising effectiveness of untrained networks. In contrast to studies of the primate visual system, classification-trained networks appear to have little-to-no benefit over untrained convolutional networks when predicting the visual cortex representations of mice (Cadena et al., 2019). Thus, when considering biological vision more broadly, there may still be much to learn from architectural principles alone, independent of task-based pre-training (Shi et al., 2019).

Our findings suggest potential new directions for theoretical research on random-feature computation. Several lines of theoretical work explore networks whose hidden layers contain random features (Chang & Futagami, 2019; Jaeger, 2001; Mei et al., 2021; Mei & Montanari, 2020; Rahimi & Recht, 2007). Like these approaches, we investigated networks with wide random-feature layers, and we only fit the output layer. However, our work differs in several important ways from these previous theoretical studies. First, classical random feature models typically use simpler fully connected or recurrent architectures, often with a single hidden layer (though they can also be stacked on top of more complex feature extraction methods, such as convolutional networks (Tong & Tanaka, 2018)). In contrast, our work specifically examines how wide random feature layers within different deep architectures (CNNs, transformers, fully connected networks) affect performance. Our finding that CNNs show markedly better performance highlights how architectural inductive biases interact with random features in ways not typically studied in the random feature literature. Second, our work in inspired by neurobiological theories and focuses specifically on the ability of random convolutional networks to predict cortical image representations. Third, our results suggest that the spatial locality and hierarchical structure of CNNs create emergent representational properties that are not present in simpler random feature models. Future work exploring why CNNs show such different behavior from other architectures in the high-dimensional regime could advance both machine learning theory and our understanding of biological vision. Furthermore, it is known that certain architectural components, including activation functions, layer normalizations, and residual connections, can bias networks to learn low complexity functions that tend to be well-aligned with real-world data (Teney et al., 2024). Our findings suggest that another promising direction for future work is to explore whether convolutional architectures are biased to learn functions whose complexity is well-aligned with mammalian vision.

Our findings about dimensionality expansion may also have interesting connections to theoretical work on neural networks in the infinite-width limit, particularly the neural tangent kernel (NTK) (Jacot et al., 2018). In the NTK regime (i.e., very wide networks), weights only need to change slightly during training, indicating that the network was already close to a solution at initialization. This phenomenon has conceptual similarities to our findings, which show that certain wide convolutional architectures appear close to a brain-aligned representation at initialization. However, there are important differences between our work and NTK theory. First, our work examines untrained networks with finite (though large) widths, and we find that performance remains high even when the width of the network is dramatically reduced through PCA. This contrasts with the infinite-width setting of the NTK. Second, our networks achieve strong performance when training only the output regression layer and keeping all other layers frozen, whereas NTK theory describes the dynamics of gradient-based learning with weight updates in all layers. Thus, despite some intriguing parallels, our findings cannot be readily explained by NTK theory, and an exciting open question is whether future extensions of NTK theory could shed light on the finite-width setting and architecture-dependent effects reported here.

We observed a notable difference in the encoding performance of our untrained CNN in the monkey data compared with the human data. While the untrained CNN rivaled AlexNet in monkey V4 and IT, it had a substantial performance gap relative to AlexNet in the human data, and this performance gap increased at higher levels of the ventral stream. This difference between the monkey and human data may be caused by several factors, such as differences in the stimulus sets and experimental tasks or representational differences across species, with human vision potentially placing a stronger emphasis on semantic associations (Bonner & Epstein, 2021; Doerig et al., 2022; Wang et al., 2023). However, another possible explanation is the differential contribution of feedback processes in these two datasets. In the monkey electrophysiology study, the images were presented for 100 ms to elicit purely perceptual processes, and neural responses were taken from a small temporal window immediately after stimulus onset to measure feedforward activations. In contrast, in the human fMRI study, the images were shown for three seconds, and due to the low temporal resolution of fMRI, the responses reflect at least several seconds of cortical activity, which means that they have a much greater contribution from feedback and recurrent processes that may emphasize abstract semantic information.

Although other work has argued for random dimensionality expansion as a biological computation (Babadi & Sompolinsky, 2014; Cayco-Gajic & Silver, 2019), we want to emphasize that our findings do not indicate that visual cortex itself performs such a random expansion. In our work, dimensionality expansion serves not as a proposed computational mechanism but rather as a means of discovering brain-relevant representations, just as others have used supervised pre-training and backpropagation. However, note that there is a key distinction between the dimensionality-expansion approach used here and the classic supervised pre-training approach. Namely, while pre-training involves a nonlinear warping of a network’s representations to achieve brain-alignment, our approach allows for brain-aligned latent dimensions to be discovered through linear regression, without the need for nonlinear fitting. Intriguingly, the process of distilling brain-relevant representation from a random, high-dimensional feature space has potential parallels with the neural pruning that occurs in visual cortex over the course of development (Sakai, 2020). Thus, an exciting direction for future work is to consider whether pruning in high-dimensional feature spaces could be incorporated into computational models of learning in human vision. Another potential direction is to consider whether our architecture could be combined with a small amount of pre-training to yield a highly performant model that is trained in a data-efficient manner.

In sum, our results show that many aspects of visual cortex representation naturally emerge in high-dimensional convolutional networks without the need for massive pre-training. Thus, contrary to the view that many diverse architectures explain visual cortex representations equally well (Conwell et al., 2022; Storrs et al., 2021), our findings show that convolutional architectures are exceptional in their ability to predict cortical representations de novo.

## METHODS

### Neural network architectures

#### Convolutional architecture

Our convolutional network, referred to as the Expansion model in this paper, is constructed by repeating a block of operations across a hierarchy of layers. Our goal was to predict image-evoked responses in higher-level visual cortex, so we constructed a deep network with five layers. The same base architecture was used for all analyses, with changes made to the dimensionality of the final layer. Each layer of the model consists of the following operations: convolution, ReLU, and average pooling. In layer 1, we used pre-defined wavelets with a curvature parameter. The use of these curved wavelets allows the model to detect both low-level straight contours and mid-level curved contours, and it was inspired by previous work that used similar wavelets to model mid-level representations in the visual cortices of humans and macaques (Bruna & Mallat, 2012; Pogoncheff et al., 2023; Yue et al., 2014, 2020). We constructed these wavelets using 3 degrees of curvature and 12 orientations, resulting in 36 filters, which were applied separately to each RGB channel to yield 108 output channels. We note that model performance was not strongly contingent on the specific number of curvature degrees and orientations. Convolution layers 2-5 consist of randomly initialized filters constructed with kaiming uniform initialization in PyTorch. We used a fixed number of filters for layers 2-4 and a varied number of filters in layer 5. The total number of parameters in the Expansion models ranges from approximately 215 million for the smallest model to 1.5 billion for the largest. For comparison, AlexNet has about 61 million trainable parameters. However, it is important to note that most of the parameters in our Expansion model are not actually trained (we only train a linear regression at the output layer), whereas all the parameters in supervised AlexNet are trained. The maximum output dimensionality for layer 5 differed across the two neural datasets due to memory limitations during the model-fitting procedure; the smaller size of the macaque electrophysiology data enabled us to explore the effects of expansion up to 10^6^ features, whereas with the larger fMRI data, we could only expand up to 10^5^ features. Similarly, to avoid memory issues during principal component analysis, a model with 10^5^ features was used for image classification. Further details about this architecture are listed in Supplementary Table 1.

#### Fully connected architecture

We created a five-layer fully connected network in which the number of units in each layer matched the number of channels in the corresponding layer of the Expansion model. As in the Expansion model, the architecture was fixed for layers 1-4, and we varied the number of features in the final layer. Further details about this architecture are listed in Supplementary Table 2.

#### Transformer architecture

We used an untrained Vision Transformer (ViT) based on the architectural specifications of the “vit_base_patch16_224” model from the timm Python library (*Pytorch Image Models (Timm) | Timmdocs*, n.d.). To expand the network’s dimensionality in a manner consistent with the other architectures, we focus on the final encoder block. We insert a linear projection after layer 11 to map the token embeddings from 768 dimensions to a new dimensionality N∈{12,60,600,6000} (see Supplementary Table 3). This projected output is then passed to the final transformer block, which now operates on N-dimensional embeddings instead of the original 768. As a result, the output of the final block has shape 196×N, where 196 is the number of tokens minus the CLS (classification token) in the original ViT (corresponding to a 224×224 image with 16×16 patches). We report this 196×N as the feature dimensionality of the transformer in all main analyses. The values of N were chosen to match the order of magnitude of feature dimensions used in our Expansion and fully connected models. Across these models, we did not perform an explicit space-versus-feature factorization. Instead, we select features such that the total feature dimensionality—computed as the product of spatial and feature dimensions—falls within a similar range across models.

### Neural datasets

Human fMRI data were obtained from a publicly available dataset, called the Natural Scenes Dataset (NSD) (Allen et al., 2022), that contains fMRI responses to 73,000 natural images from the Microsoft Common Objects in Context (COCO) database (Lin et al., 2015). These fMRI data were collected at ultra-high-field strength with high spatial resolution (7T field strength, 1.6-s TR, 1.8mm^3^ voxel size). Each image was shown for three seconds, and participants were instructed to maintain fixation. We used the NSD single-trial betas, preprocessed in 1.8-mm volume space and denoised using the GLMdenoise technique (version 3; “betas_fithrf_GLMdenoise_RR”) (Kay et al., 2013). We converted the betas to z-scores within each scanning session and computed the average betas for each stimulus across repetitions. Each participant viewed up to 10,000 images, of which 1,000 were shown to all participants and 9,000 were unique to each participant. We examined responses to all images shown to all 8 participants, and we focused on voxels from the ventral visual pathway using regions of interest defined in the NSD data as early, midventral, and ventral visual streams. These regions were predefined by the original authors as part of the “streams” ROIs. In Extended Data Figure 1, we also examined fMRI data from the large-scale THINGS dataset, which contains fMRI responses to 8,640 object images in three subjects (Hebart et al., 2023).

Macaque electrophysiology data were obtained from a publicly available dataset containing single-unit recordings from two macaques implanted with arrays of electrodes in V4 and IT (Majaj et al., 2015). Neural responses were recorded from a total of 88 sites in V4 and 168 in IT. For each site, the average response was obtained between 70 and 170 ms after stimulus presentation. The stimuli in this dataset consisted of 3200 artificially generated gray-scale images composed from 64 objects belonging to 8 categories. The objects were positioned on top of natural scenes at various locations, orientations, and scales. All images were seen by both animals.

### Encoding models

We used linear regression with L2 regularization to map model activations to unit and voxel responses. For these analyses, the data were split into sets of training and testing images. In the human fMRI data, the training set for each participant consisted of all images that were uniquely seen by that participant, while the test set consisted of the images that were seen by all participants. In the monkey electrophysiology data and THINGS fMRI data, the training and test sets were selected randomly with an 80:20 split.

To identify the best mapping for each model, we first found the optimal L2 penalty for each participant and each region through leave-one-out cross validation on the training images. We tested 10 L2 penalty values logarithmically spaced using base 10, ranging from 1 to 10^10^. We selected the optimal penalty that results in the best average score across all units or voxels, and we used the optimal penalty to estimate regression coefficients on the entire training set. The estimated coefficients were then applied to the model activations for the held-out test images to generate predicted neural responses.

To assess the accuracy of these predictions, we calculated the Pearson correlation coefficient (r value) of the predicted and actual responses for each unit and voxel. We computed bootstrap distributions of these accuracy scores over 1,000 iterations in which the test images were resampled with replacement. The encoding performance score for each region was obtained by taking the average of these distributions over all units or voxels and all participants. For the untrained networks, image features were extracted from the last layer and used as regressors. For AlexNet, we first extracted features from all the convolutional layers of the model. The number of features from each layer is equal to [channels x height x width]. For instance, the final convolutional layer (Conv5) produces 256 feature maps of size 6×6, resulting in a total of 9216 features. The top-performing layer for each region was then identified through cross-validation on the training images, following the same procedure used to determine the optimal L2 penalty.

For analyses involving PCA, we learned PCs of network activations on the training images, and we projected the activations for both the training and test images onto these PCs. We then performed encoding model analyses on these reduced network representations using the procedure described above.

We computed estimates of upper bounds for the noise ceilings in each ROI, which indicate the maximum possible correlation that one could expect to find given the internal reliability of the neural responses. Specifically, we first computed Spearman-Brown adjusted correlations of the unit-wise and voxel-wise responses across splits of repeated stimulus presentations. In the monkey data, we used a random 50:50 split of the stimulus repetitions, and in the human data, which had only three repetitions at most, we computed the mean correlation across all possible pairwise comparisons of repetitions. We then computed as estimate of the upper bound based on the correlation attenuation formula (Spearman, 1904). Specifically, for a correlation between two variables, x and y, the upper bound is equal to the square root of the product of the internal reliabilities of x and y. Because the neural networks are deterministic, their reliabilities are equal to 1, which means that the upper bound for the encoding score is equal to the square root of the reliability of the neural data.

### Image classification

Image features were extracted from the last convolution layer of each model using the Places365-Standard dataset (Zhou et al., 2018), which contains 365 categories of scenes. To remove any effect of feature dimensionality on classification, we learned the first 1,000 PCs from each model on a random subset of 100 images from each category in the training set (for a total of 36,500 images), and projected model features from the entire validation set onto these learned PCs. Each model’s PCs were then used for classification on the validation set, which contains 100 images per category. Classification accuracy was determined through 5-fold cross-validation using sklearn’s K-Fold cross-validator, which ensures a consistent sample distribution across each class. Each fold was split into an 80:20 distribution, resulting in 80 train images and 20 test images per class. Using sklearn’s Logistic Regression classifier, images were classified into categories in a 365-way classification, with chance performance at 1/365 or 0.0027.

## DATA AVAILABILITY

The Natural Scenes Dataset is available here: https://naturalscenesdataset.org/.

The THINGS fMRI dataset is available here: https://openneuro.org/datasets/ds004192

The monkey electrophysiology dataset is available as part of the Brain-score GitHub package: https://github.com/brain-score.

The Places dataset is available here: http://places2.csail.mit.edu/download.html.

## CODE AVAILABILITY

The Expansion model, as well as code for all analyses is available on GitHub: https://github.com/akazemian/untrained_models_of_visual_cortex.

## SUPPLEMENT

**Supplementary Table 1:**
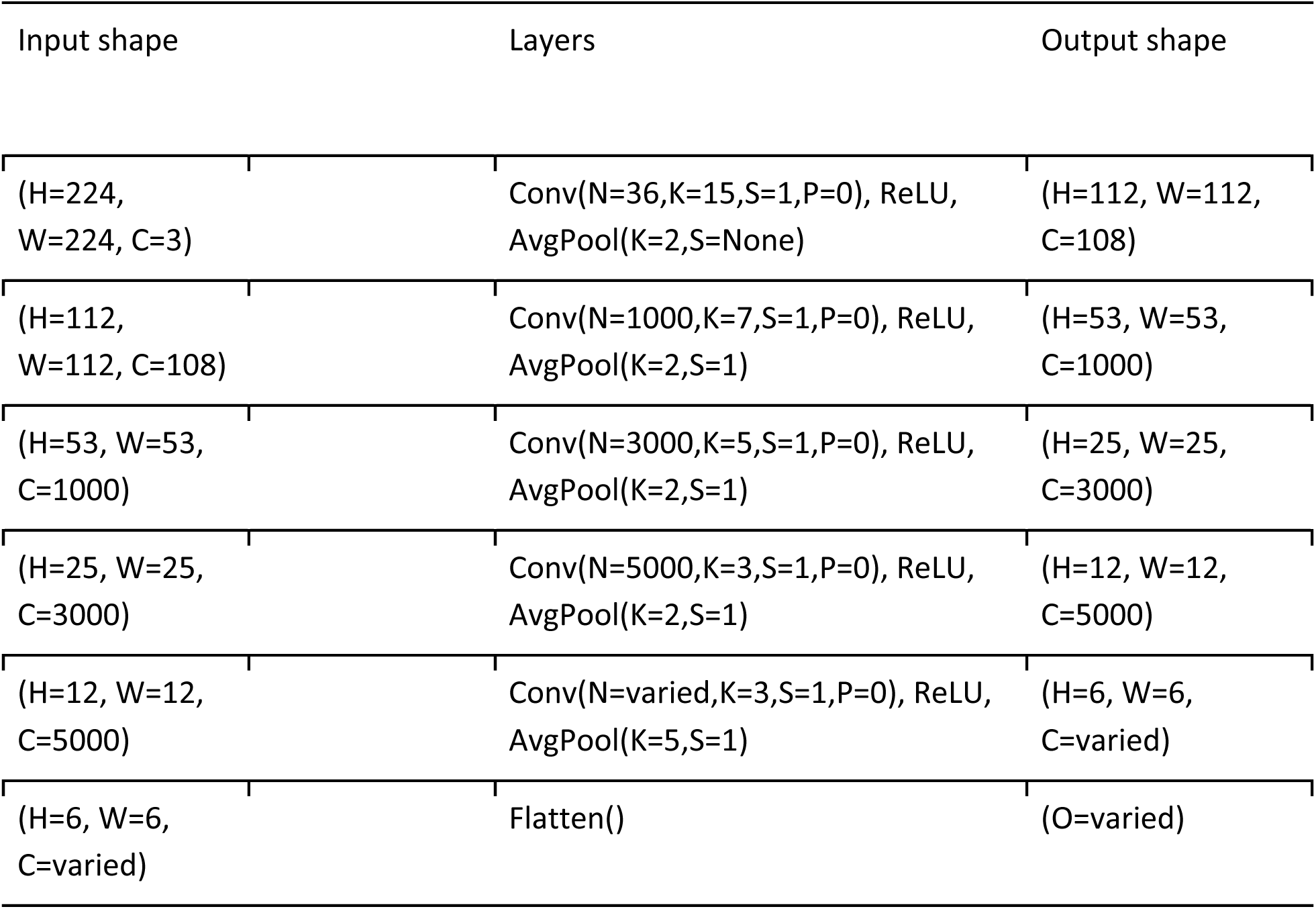
Expansion model architecture. The number of filters in the last layer is varied to study the effect of feature expansion. H, W and C stand for height, width and number of channels respectively. In each layer, N, K, S and P stand for the number of convolution filters, filter size, stride, and padding.

| Input shape | Layers | Output shape |
| --- | --- | --- |
| (H=224, W=224, C=3) | Conv(N=36,K=15,S=1,P=0), ReLU,<br>AvgPool(K=2,S=None) | (H=112, W=112, C=108) |
| (H=112, W=112, C=108) | Conv(N=1000,K=7,S=1,P=0), ReLU,<br>AvgPool(K=2,S=1) | (H=53, W=53, C=1000) |
| (H=53, W=53, C=1000) | Conv(N=3000,K=5,S=1,P=0), ReLU,<br>AvgPool(K=2,S=1) | (H=25, W=25, C=3000) |
| (H=25, W=25, C=3000) | Conv(N=5000,K=3,S=1,P=0), ReLU,<br>AvgPool(K=2,S=1) | (H=12, W=12, C=5000) |
| (H=12, W=12, C=5000) | Conv(N=varied,K=3,S=1,P=0), ReLU,<br>AvgPool(K=5,S=1) | (H=6, W=6, C=varied) |
| (H=6, W=6, C=varied) | Flatten() | (O=varied) |

**Supplementary Table 2:**
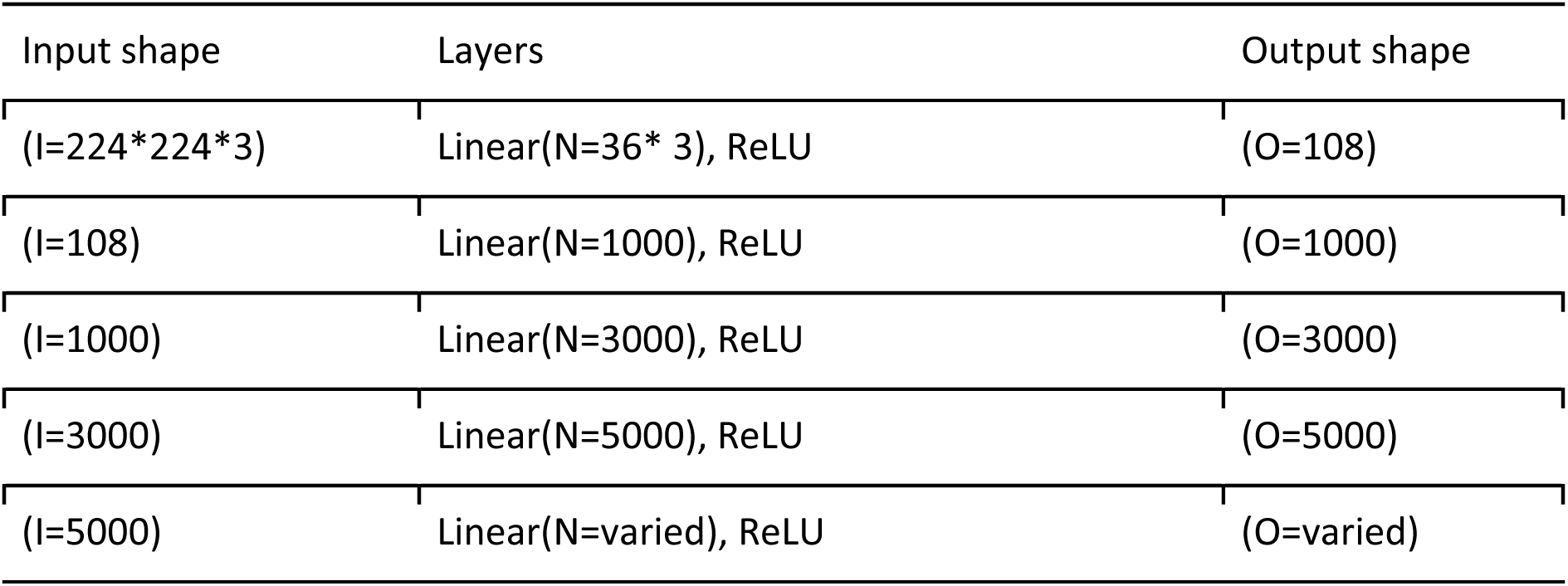
Fully connected architecture. The number of filters in the last layer is varied for comparison against the Expansion model. I and O stand for input and output size, and N stands for the size of the linear layer.

**Supplementary Table 3:** Untrained transformer architecture. The number of embeddings in the last block is varied for comparison against the Expansion model. For simplicity, this table shows only the operations that are different from a classic ViT architecture. Therefore, the input has <u>the shape of the output from block 11. N stands for the size of the linear layer.</u>

| Input shape | Layers | Output shape |
| --- | --- | --- |
| (196, 768) | Linear(N) | (196, N=Varied) |
| (196, N) | ViT Block with embedding dimension = N | (196, N) |

## EXTENDED DATA FIGURES

**Extended Data Figure 1.**
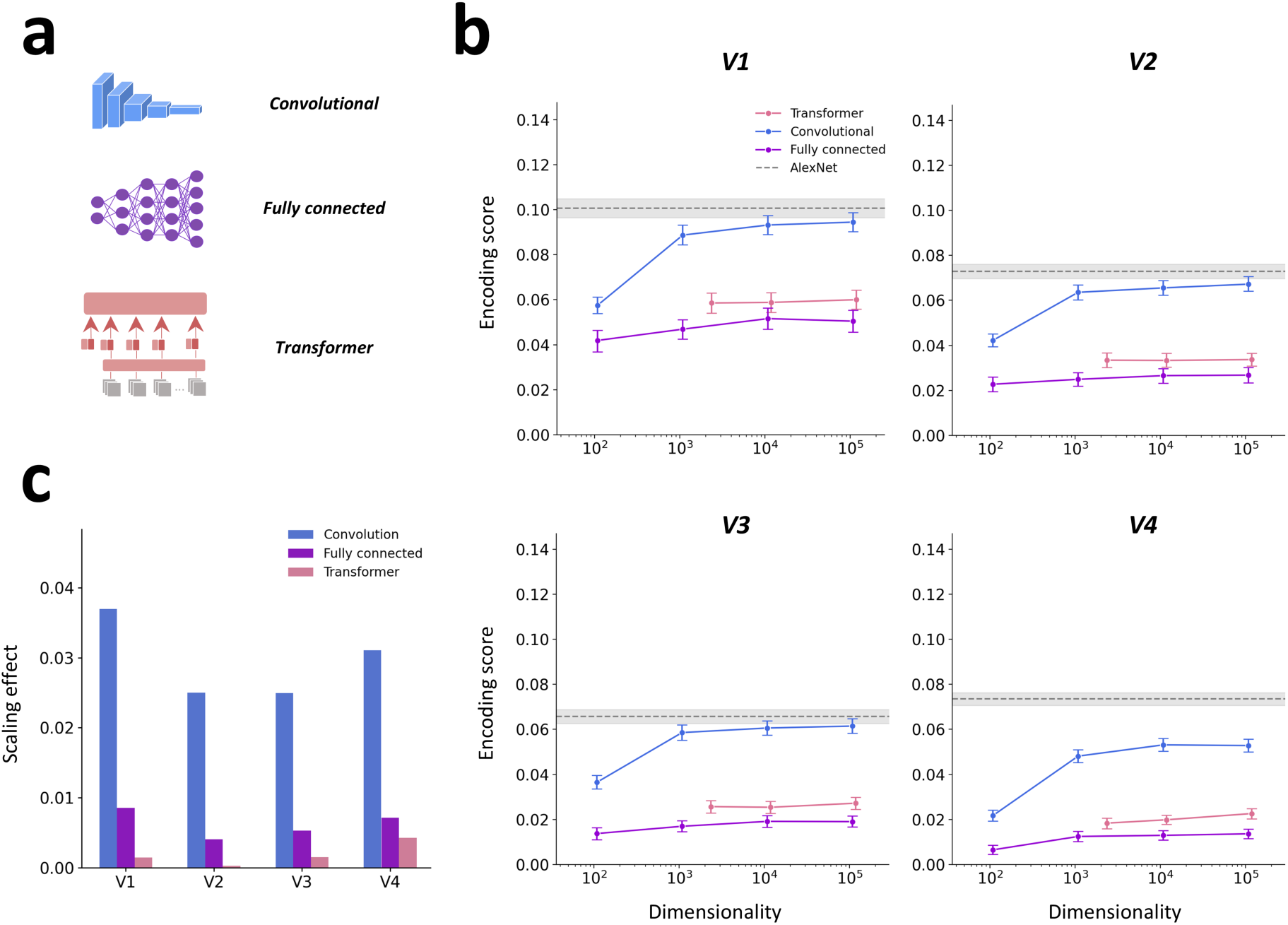
Dimensionality expansion strongly improves the encoding performance of convolutional neural networks in another large-scale fMRI dataset. **a)** The effects of dimensionality expansion were examined for a convolutional neural network, a deep fully connected network, and a vision transformer. **b)** These plots illustrate the encoding performance for all three architectures as a function of dimensionality expansion. There was no pre-training for these networks. Encoding performance was evaluated for regions V1 to V4 from the large-scale THINGS fMRI dataset. The x-axis plots the number of random features in the output layer, and the y-axis shows the encoding score for predicting image-evoked cortical responses. The gray dashed line indicates the performance of the best performing convolutional layer of pre-trained AlexNet. The convolutional architecture without pre-training attained large performance gains as a function of dimensionality expansion. In contrast, the other two architectures showed much less improvement as the number of output features was expanded. The encoding performance plots show the mean performance across voxels from all participants, and the error bars denote 97.5% confidence intervals from 1,000 bootstrap samples. **c)** A summary statistic for the effect of dimensionality expansion was calculated as the difference in performance between the highest- and lowest-dimensional version of each architecture. The performance of the convolutional model was strongly modulated by the dimensionality of its random feature space. In contrast, the effect of dimensionality expansions was much weaker in the other two architectures.

**Extended Data Figure 2.**
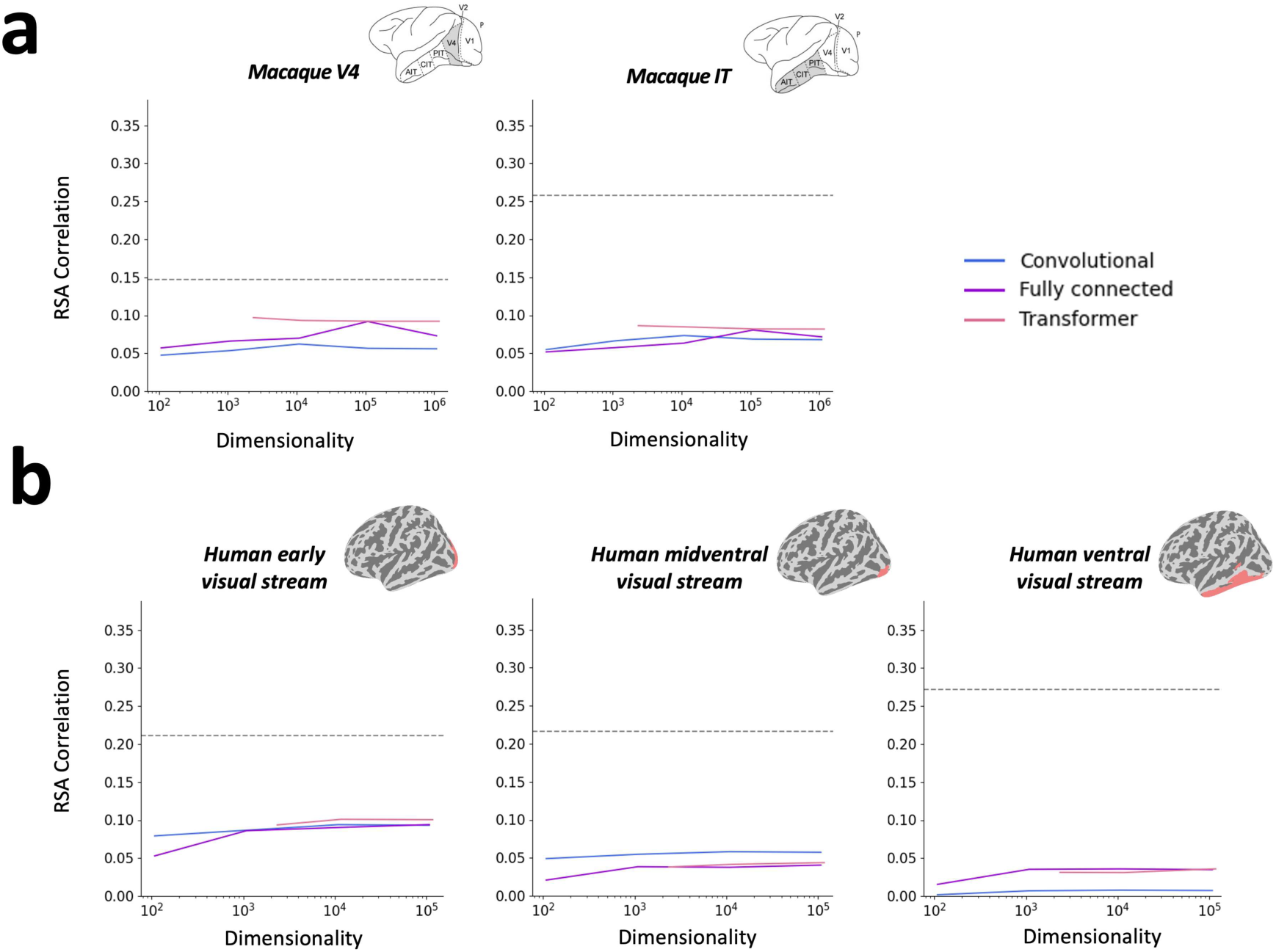
The benefits of dimensionality expansion depend on linear reweighting. **a)** The effects of dimensionality expansion were examined for a convolutional neural network, a deep fully connected network, and a vision transformer. **b)** These plots illustrate representational similarity analysis (RSA) scores for all three architectures as a function of dimensionality expansion. There was no pre-training for these networks. RSA scores were evaluated for regions V4 and IT in the monkey electrophysiology data. The x-axis plots the number of random features in the output layer, and the y-axis shows the RSA score. The gray dashed line indicates the performance of the best performing convolutional layer of pre-trained AlexNet. As expected, the RSA scores do not systematically increase as a function of dimensionality expansion. This contrasts with the effect of dimensionality expansion observed for the encoding scores in Figure 2, and it demonstrates that a linear reweighting procedure is needed to yield performance gains from dimensionality expansion in untrained networks. The encoding performance plots in this panel and in panel c show the mean performance across units/voxels from all participants, and the error bars denote 97.5% confidence intervals from 1,000 bootstrap samples. **c)** These plots show the same analyses as in panel b but for regions along the ventral stream in the human fMRI data.

**Extended Data Figure 3.**
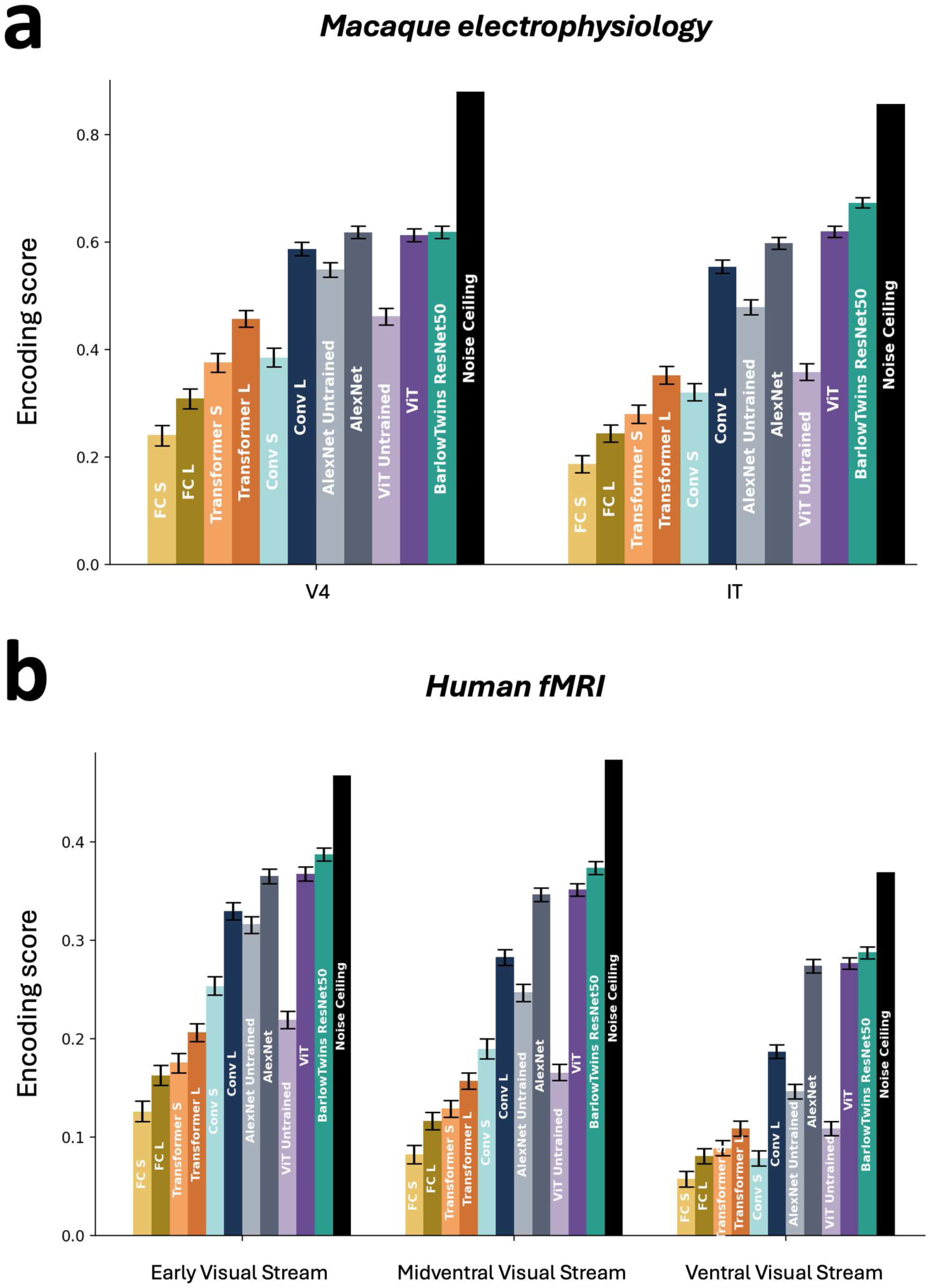
Comparisons with other trained deep neural network architectures. **a)** These plots show the encoding performance of state-of-the-art trained architectures for comparison with the untrained networks and trained AlexNet. For each untrained architecture, these plots show performance at both the smallest and largest levels of dimensionality. In addition to AlexNet, these plots include more modern architectures, specifically ViT and BarlowTwins, pre-trained on ImageNet. Encoding performance was evaluated for regions V4 and IT in the monkey electrophysiology data. The encoding performance plots in this panel and in panel b show the mean performance across units/voxels from all participants, and the error bars denote 97.5% confidence intervals from 1,000 bootstrap samples. These plots all show the mean noise ceiling across units/voxels. **b)** These plots show the same analyses as in panel a but for regions along the ventral stream in the human fMRI data.

**Extended Data Figure 4.**
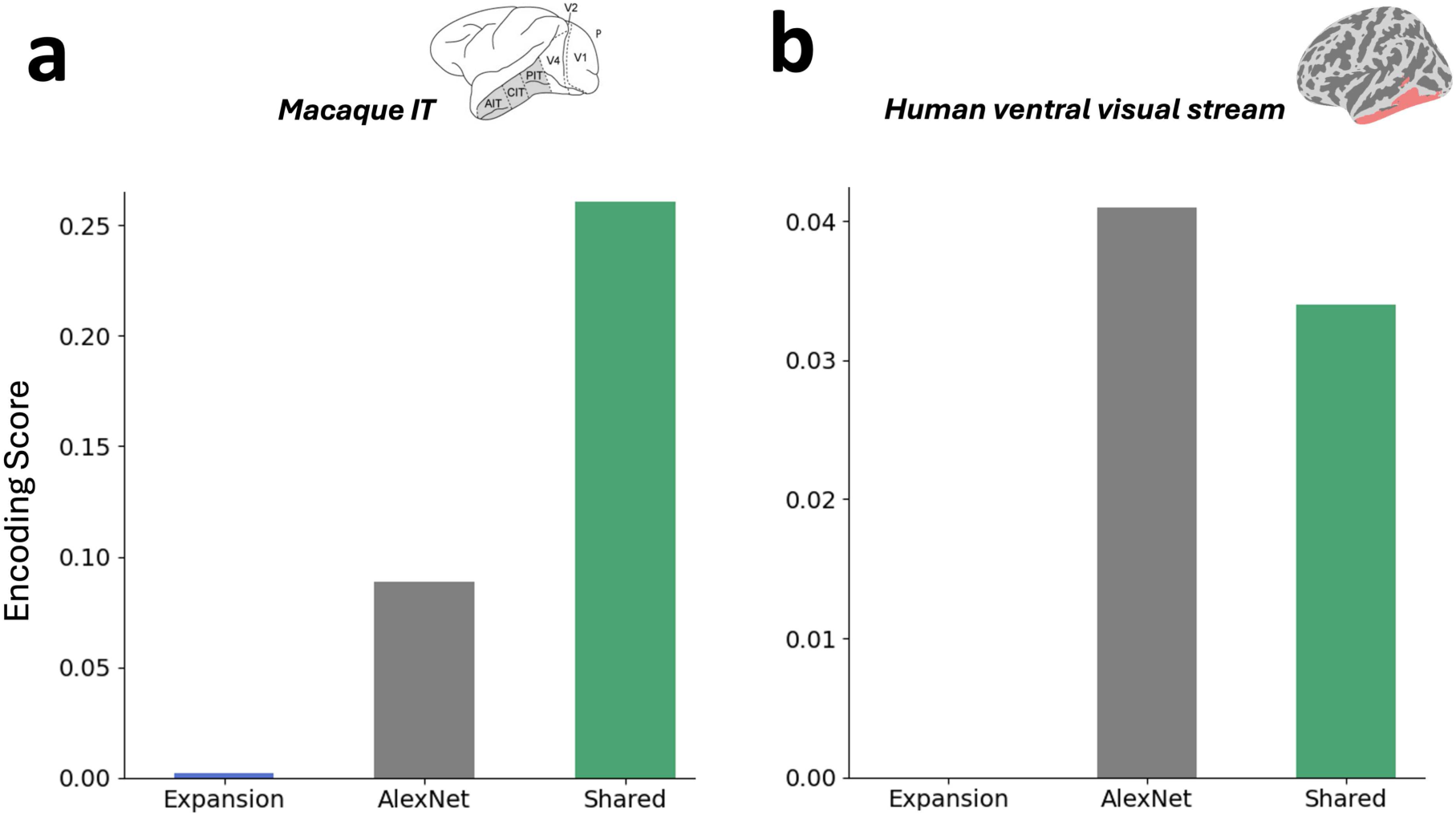
Variance partitioning shows that the Expansion model explains the same variance as trained AlexNet. Variance partitioning was used to determine whether the Expansion model explains the same variance in cortical responses as trained AlexNet. These analyses were performed using the largest convolutional model, and they show that in both the monkey electrophysiology data (left) and human fMRI data (right) the explained variance of the Expansion model is fully shared with AlexNet. As expected, AlexNet explains additional unique variance that is not shared with the Expansion model.

**Extended Data Figure 5.**
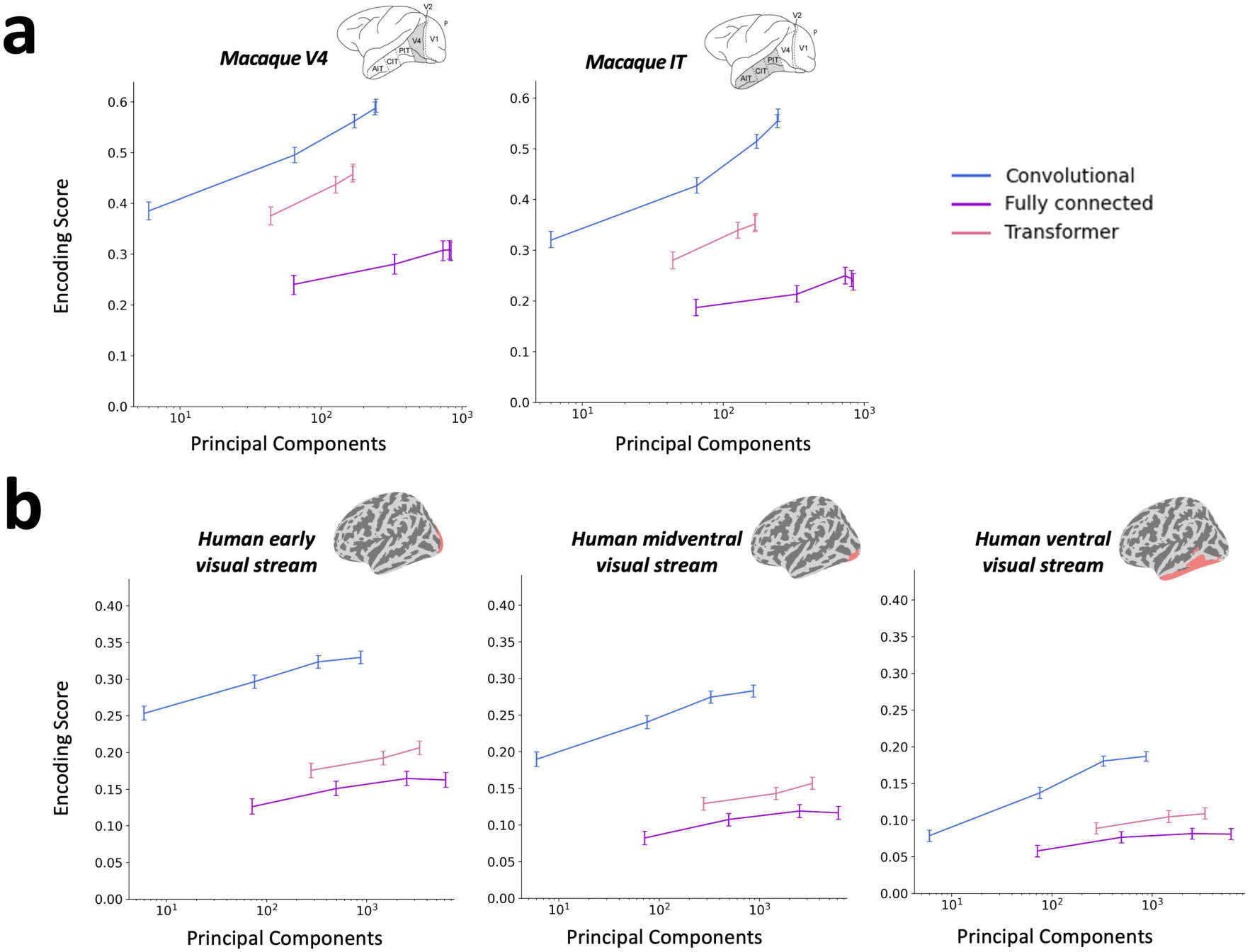
Improvements in encoding performance are linked to increases in latent dimensionality. **a)** These plots illustrate the encoding performance of the networks from Figure 2 as a function of their latent dimensionality. Encoding performance was evaluated for regions V4 and IT in the monkey electrophysiology data. Plotting conventions are the same as in Figure 2, except that here the x-axis plots the number of principal components that account for 85% of variance in the output layer. The results show that improvements in encoding performance are closely linked to increases in the latent dimensionality of a network’s natural image representations. However, note that measures of latent dimensionality are architecture-dependent, and thus different architectures with the same level of latent dimensionality can nonetheless differ in encoding performance. **b)** These plots show the same analyses as in panel a but for regions along the ventral stream in the human fMRI data.

**Extended Data Figure 6.**
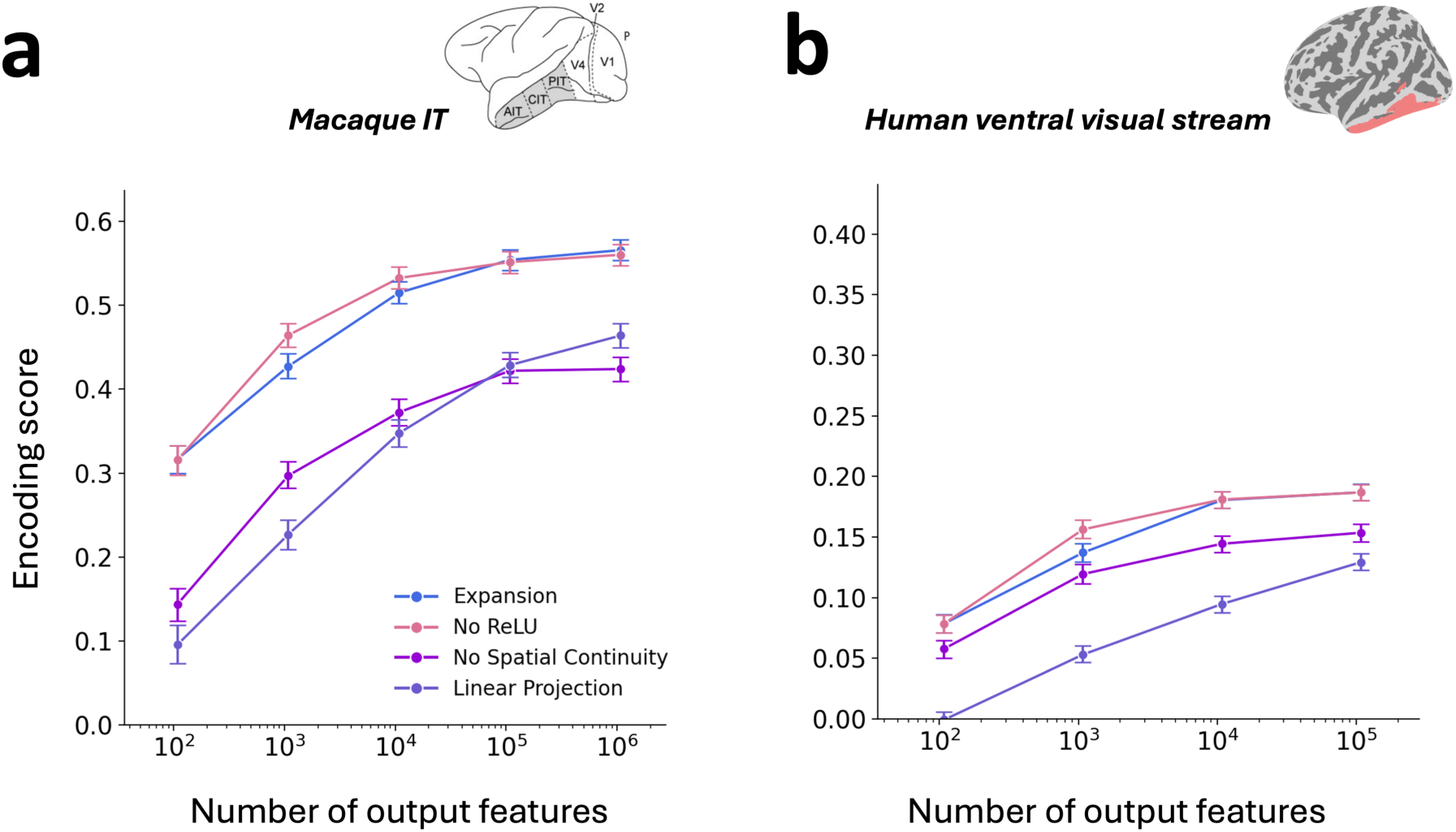
Analysis of architectural components in the final layer of the convolutional network. **a-b)** These plots show the performance of the largest untrained convolutional network after altering key architectural components in the final layer only. Panel a shows results for macaque IT, and panel b shows results for the human high-level ventral stream. These plots show that ablating the nonlinearity in the final layer has little effect, which means that the crucial nonlinear operations are those that occur in earlier layers. They also show that removing the spatial locality of the convolutional filters in the final layer results in an overall decrease in encoding performance, demonstrating that even in the highest layer of the network it is beneficial to compute spatially local representations. Plotting conventions are the same as in Figure 2.

**Extended Data Figure 7.**
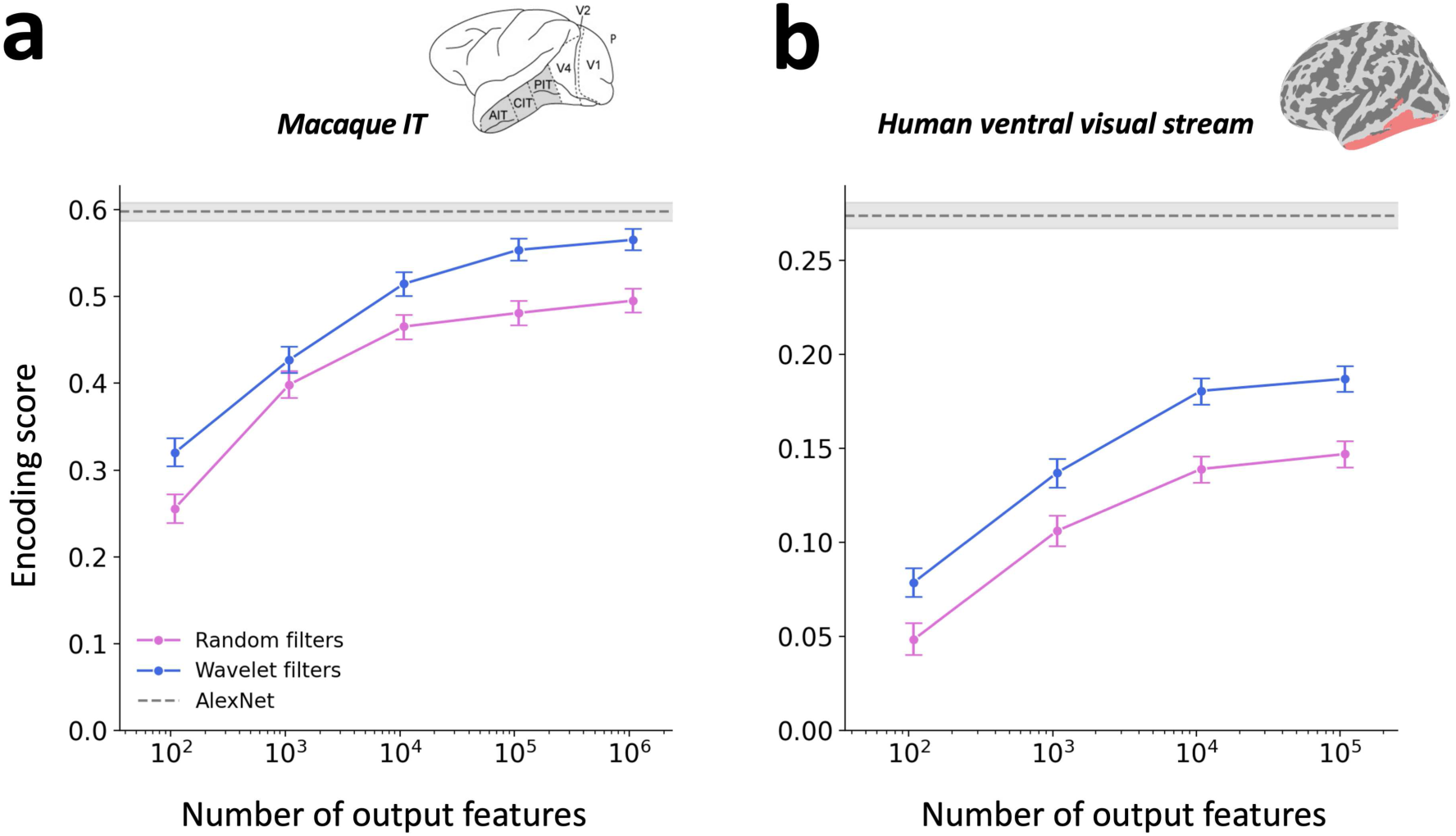
Effect of pre-defined wavelets in the first layer of the untrained convolutional neural network. These plots show the effect of using pre-defined wavelets in the first layer of the untrained convolutional network. Encoding performance is shown for macaque IT (left) and the human high-level ventral stream (right). For comparison, we examined a model with 3,000 randomly initialized filters in layer 1 (the number of random filters was maximized within computational memory limits). As in Figure 2, these plots show how encoding performance changes as a function of dimensionality expansion in the final convolutional layer. The x-axis plots the number of random features in the output layer, and the y-axis shows the encoding score for predicting image-evoked cortical responses. The gray dashed line indicates the performance of the best-performing convolutional layer of pre-trained AlexNet. There is a small, but consistent drop in performance for the fully random model across both datasets, demonstrating that overall, the network benefits from the implementation of pre-defined wavelets in its first layer.

**Extended Data Figure 8.**
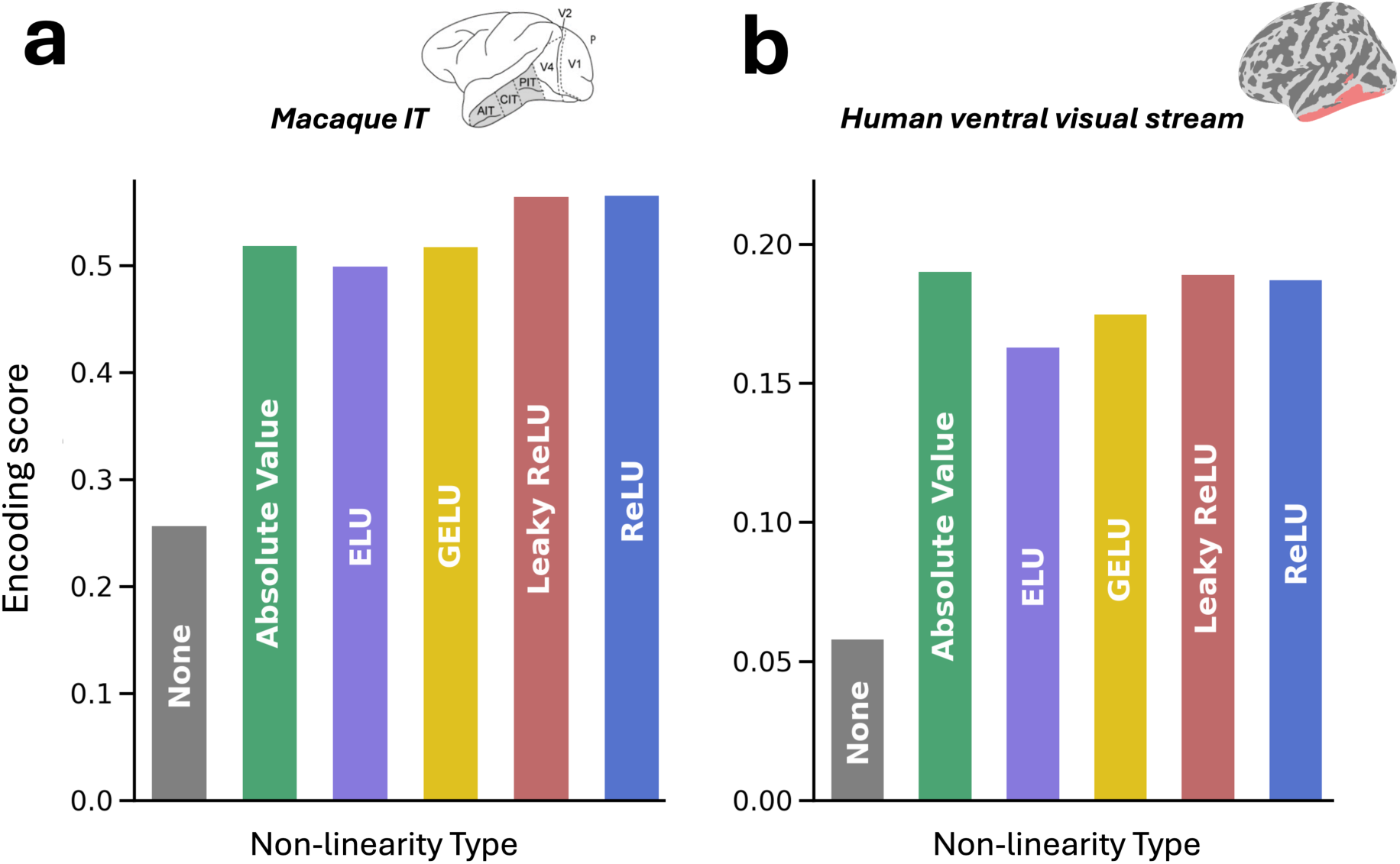
Different activation functions yield similar encoding performance for the untrained convolutional neural network. The effects of using different nonlinear activation functions in our untrained convolutional network were explored for monkey IT (left) and the human high-level ventral stream (right). These plots illustrate encoding performance for models with different nonlinear activation functions with otherwise identical architectures, all containing 10^5^ features in their output layer. For comparison, this plot also includes a network without any nonlinear activation functions (the Linear model). The y-axis shows the encoding score for predicting image-evoked cortical responses. The results demonstrate that the inclusion of nonlinearities is critical, but various types of nonlinearities yield similar levels of performance. ReLU = rectified linear unit, GELU = Gaussian error linear unit, ELU = exponential linear unit

**Extended Data Figure 9.**
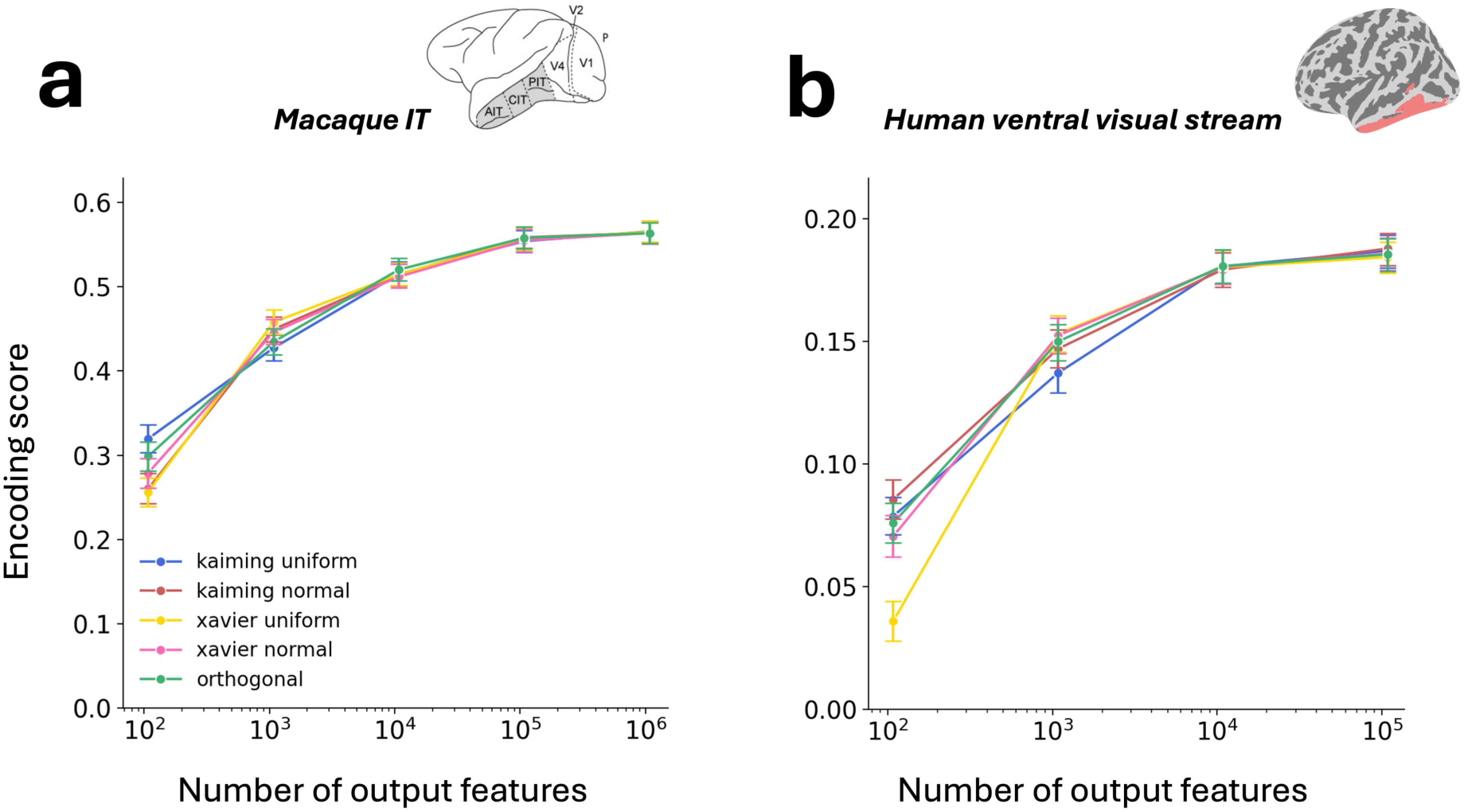
Different random initialization methods yield similar encoding performance for the untrained convolutional neural network. The effects of initializing the random features of our untrained convolutional network using different methods were explored for monkey IT (left) and human ventral visual stream (right). These plots illustrate encoding performance for identical architectures with different random initialization types. As in Figure 2, the x-axis plots the number of random features in the output layer, and the y-axis shows the encoding score for predicting image-evoked cortical responses. There is variation in encoding performance for different initialization methods in models with low dimensionality (on the left side of the x-axis). However, at higher levels of dimensionality, these performance differences diminish. This indicates that the type of initialization has minimal impact on encoding performance in the presence of model expansion.

## Notes

https://github.com/akazemian/untrained_models_of_visual_cortex

### Competing Interest Statement

The authors have declared no competing interest.

### Summary of Updates

Updated results and a more extensive list of supplementary figures.

## REFERENCES

Agrawal, P., Stansbury, D., Malik, J., & Gallant, J. L. (2014). Pixels to Voxels: Modeling Visual Representation in the Human Brain. arXiv:1407.5104 [Cs, q-Bio]. http://arxiv.org/abs/1407.5104

Allen, E. J., St-Yves, G., Wu, Y., Breedlove, J. L., Prince, J. S., Dowdle, L. T., Nau, M., Caron, B., Pestilli, F., Charest, I., Hutchinson, J. B., Naselaris, T., & Kay, K. (2022). A massive 7T fMRI dataset to bridge cognitive neuroscience and artificial intelligence. Nature Neuroscience, 25(1), 116–126. 10.1038/s41593-021-00962-x

Babadi, B., & Sompolinsky, H. (2014). Sparseness and Expansion in Sensory Representations. Neuron, 83(5), 1213–1226. 10.1016/j.neuron.2014.07.035

Baek, S., Song, M., Jang, J., Kim, G., & Paik, S.-B. (2021). Face detection in untrained deep neural networks. Nature Communications, 12(1), 7328. 10.1038/s41467-021-27606-9

Bonner, M. F., & Epstein, R. A. (2021). Object representations in the human brain reflect the co-occurrence statistics of vision and language. Nature Communications, 12(1), 4081. 10.1038/s41467-021-24368-2

Bruna, J., & Mallat, S. (2012). *Invariant Scattering Convolution Networks* (arXiv:1203.1513). arXiv. 10.48550/arXiv.1203.1513

Cadena, S. A., Denfield, G. H., Walker, E. Y., Gatys, L. A., Tolias, A. S., Bethge, M., & Ecker, A. S. (2019). Deep convolutional models improve predictions of macaque V1 responses to natural images. PLOS Computational Biology, 15(4), e1006897. 10.1371/journal.pcbi.1006897

Cao, R., & Yamins, D. (2021a). *Explanatory models in neuroscience: Part 1 -- taking mechanistic abstraction seriously* (arXiv:2104.01490). arXiv. 10.48550/arXiv.2104.01490

Cao, R., & Yamins, D. (2021b). *Explanatory models in neuroscience: Part 2 -- constraint-based intelligibility* (arXiv:2104.01489). arXiv. 10.48550/arXiv.2104.01489

Cao, Y.-H., & Wu, J. (2021). *A Random CNN Sees Objects: One Inductive Bias of CNN and Its Applications* (arXiv:2106.09259). arXiv. 10.48550/arXiv.2106.09259

Carandini, M., Demb, J. B., Mante, V., Tolhurst, D. J., Dan, Y., Olshausen, B. A., Gallant, J. L., & Rust, N. C. (2005). Do We Know What the Early Visual System Does? Journal of Neuroscience, 25(46), 10577–10597. 10.1523/JNEUROSCI.3726-05.2005

Casper, S., Boix, X., D’Amario, V., Guo, L., Schrimpf, M., Vinken, K., & Kreiman, G. (2021). *Frivolous Units: Wider Networks Are Not Really That Wide* (arXiv:1912.04783). arXiv. 10.48550/arXiv.1912.04783

Cayco-Gajic, N. A., & Silver, R. A. (2019). Re-evaluating Circuit Mechanisms Underlying Pattern Separation. Neuron, 101(4), 584–602. 10.1016/j.neuron.2019.01.044

Chang, H., & Futagami, K. (2019). *Reinforcement Learning with Convolutional Reservoir Computing* (arXiv:1912.04161). arXiv. 10.48550/arXiv.1912.04161

Chen, Z., & Bonner, M. F. (2025). Universal dimensions of visual representation. Science Advances, 11(27), eadw7697.

Conwell, C., Prince, J. S., Alvarez, G. A., & Konkle, T. (2022). *Large-Scale Benchmarking of Diverse Artificial Vision Models in Prediction of 7T Human Neuroimaging Data* [Preprint]. Neuroscience. 10.1101/2022.03.28.485868

Cordonnier, J.-B., Loukas, A., & Jaggi, M. (2020). *On the Relationship between Self-Attention and Convolutional Layers* (arXiv:1911.03584). arXiv. http://arxiv.org/abs/1911.03584

Doerig, A., Kietzmann, T. C., Allen, E., Wu, Y., Naselaris, T., Kay, K., & Charest, I. (2022). *Semantic scene descriptions as an objective of human vision* (arXiv:2209.11737). arXiv. 10.48550/arXiv.2209.11737

Dosovitskiy, A., Beyer, L., Kolesnikov, A., Weissenborn, D., Zhai, X., Unterthiner, T., Dehghani, M., Minderer, M., Heigold, G., Gelly, S., Uszkoreit, J., & Houlsby, N. (2021). *An Image is Worth 16×16 Words: Transformers for Image Recognition at Scale* (arXiv:2010.11929). arXiv. 10.48550/arXiv.2010.11929

Elmoznino, E., & Bonner, M. F. (2024). High-performing neural network models of visual cortex benefit from high latent dimensionality. PLOS Computational Biology, 20(1), e1011792. 10.1371/journal.pcbi.1011792

Geiger, F., Schrimpf, M., Marques, T., & DiCarlo, J. J. (2022, January 28). Wiring Up Vision: Minimizing Supervised Synaptic Updates Needed to Produce a Primate Ventral Stream | OpenReview. https://openreview.net/forum?id=g1SzIRLQXMM

Hebart, M. N., Contier, O., Teichmann, L., Rockter, A. H., Zheng, C. Y., Kidder, A., Corriveau, A., Vaziri-Pashkam, M., & Baker, C. I. (2023). THINGS-data, a multimodal collection of large-scale datasets for investigating object representations in human brain and behavior. eLife, 12, e82580. 10.7554/eLife.82580

Ingrosso, A., & Goldt, S. (2022). Data-driven emergence of convolutional structure in neural networks. Proceedings of the National Academy of Sciences, 119(40), e2201854119. 10.1073/pnas.2201854119

Jacot, A., Gabriel, F., & Hongler, C. (2018). Neural tangent kernel: Convergence and generalization in neural networks. Advances in Neural Information Processing Systems, 31.

Jaeger, H. (2001). The “echo state” approach to analysing and training recurrent neural networks-with an erratum note. *Bonn*, Germany: German National Research Center for Information Technology Gmd Technical Report, 148(34), 13.

Jarrett, K., Kavukcuoglu, K., Ranzato, M., & LeCun, Y. (2009). What is the best multi-stage architecture for object recognition? 2146–2153.

Jones, J. P., & Palmer, L. A. (1987). The two-dimensional spatial structure of simple receptive fields in cat striate cortex. Journal of Neurophysiology, 58(6), 1187–1211. 10.1152/jn.1987.58.6.1187

Kay, K. N., Rokem, A., Winawer, J., Dougherty, R. F., & Wandell, B. A. (2013). GLMdenoise: A fast, automated technique for denoising task-based fMRI data. Frontiers in Neuroscience, 7, 247. 10.3389/fnins.2013.00247

Khaligh-Razavi, S.-M., & Kriegeskorte, N. (2014). Deep supervised, but not unsupervised, models may explain IT cortical representation. PLoS Computational Biology, 10(11), e1003915. 10.1371/journal.pcbi.1003915

Konkle, T., & Alvarez, G. A. (2022). A self-supervised domain-general learning framework for human ventral stream representation. Nature Communications, 13(1), 491. 10.1038/s41467-022-28091-4

Kriegeskorte, N. (2015). Deep Neural Networks: A New Framework for Modeling Biological Vision and Brain Information Processing. Annual Review of Vision Science, 1(1), 417–446. 10.1146/annurev-vision-082114-035447

Lin, T.-Y., Maire, M., Belongie, S., Bourdev, L., Girshick, R., Hays, J., Perona, P., Ramanan, D., Zitnick, C. L., & Dollár, P. (2015). *Microsoft COCO: Common Objects in Context* (arXiv:1405.0312). arXiv. 10.48550/arXiv.1405.0312

Majaj, N. J., Hong, H., Solomon, E. A., & DiCarlo, J. J. (2015). Simple Learned Weighted Sums of Inferior Temporal Neuronal Firing Rates Accurately Predict Human Core Object Recognition Performance. Journal of Neuroscience, 35(39), 13402–13418. 10.1523/JNEUROSCI.5181-14.2015

Mei, S., Misiakiewicz, T., & Montanari, A. (2021). *Generalization error of random features and kernel methods: Hypercontractivity and kernel matrix concentration* (arXiv:2101.10588). arXiv. 10.48550/arXiv.2101.10588

Mei, S., & Montanari, A. (2020). The generalization error of random features regression: Precise asymptotics and double descent curve. *arXiv:1908.05355 [Math*, *Stat]*. http://arxiv.org/abs/1908.05355

Movshon, J. A., Thompson, I. D., & Tolhurst, D. J. (1978). Spatial summation in the receptive fields of simple cells in the cat’s striate cortex. The Journal of Physiology, 283, 53–77.

Oliva, A., & Torralba, A. (2001). Modeling the Shape of the Scene: A Holistic Representation of the Spatial Envelope. International Journal of Computer Vision, 42(3), 145–175. 10.1023/A:1011139631724

Poggio, T., Mhaskar, H., Rosasco, L., Miranda, B., & Liao, Q. (2017). Why and when can deep-but not shallow-networks avoid the curse of dimensionality: A review. International Journal of Automation and Computing, 14(5), 503–519. 10.1007/s11633-017-1054-2

Pogoncheff, G., Granley, J., & Beyeler, M. (2023). Explaining V1 Properties with a Biologically Constrained Deep Learning Architecture. In A. Oh, T. Naumann, A. Globerson, K. Saenko, M. Hardt, & S. Levine (Eds.), Advances in Neural Information Processing Systems (Vol. 36, pp. 13908–13930). Curran Associates, Inc. https://proceedings.neurips.cc/paper_files/paper/2023/file/2d1ef4aba0503226330661d74fdb236e-Paper-Conference.pdf

Portilla, J., & Simoncelli, E. P. (2000). A Parametric Texture Model Based on Joint Statistics of Complex Wavelet Coefficients. International Journal of Computer Vision, 40(1), 49–70. 10.1023/A:1026553619983

Pytorch Image Models (timm) | timmdocs. (n.d.). Retrieved May 6, 2024, from https://timm.fast.ai/

Rahimi, A., & Recht, B. (2007). Random Features for Large-Scale Kernel Machines. In J. Platt, D. Koller, Y. Singer, & S. Roweis (Eds.), Advances in Neural Information Processing Systems (Vol. 20). Curran Associates, Inc. https://proceedings.neurips.cc/paper_files/paper/2007/file/013a006f03dbc5392effeb8f18fda755-Paper.pdf

Richards, B. A., Lillicrap, T. P., Beaudoin, P., Bengio, Y., Bogacz, R., Christensen, A., Clopath, C., Costa, R. P., de Berker, A., Ganguli, S., Gillon, C. J., Hafner, D., Kepecs, A., Kriegeskorte, N., Latham, P., Lindsay, G. W., Miller, K. D., Naud, R., Pack, C. C., … Kording, K. P. (2019). A deep learning framework for neuroscience. Nature Neuroscience, 22(11), 1761–1770. 10.1038/s41593-019-0520-2

Riesenhuber, M., & Poggio, T. (1999). Hierarchical models of object recognition in cortex. Nature Neuroscience, 2(11), Article 11. 10.1038/14819

Sakai, J. (2020). How synaptic pruning shapes neural wiring during development and, possibly, in disease. Proceedings of the National Academy of Sciences, 117(28), 16096–16099. 10.1073/pnas.2010281117

Saxe, A. M., Koh, P. W., Chen, Z., Bhand, M., Suresh, B., & Ng, A. Y. (2011). On random weights and unsupervised feature learning. 2(3), 6.

Saxe, A., Nelli, S., & Summerfield, C. (2020). If deep learning is the answer, then what is the question? 26.

Schrimpf, M., Kubilius, J., Hong, H., Majaj, N. J., Rajalingham, R., Issa, E. B., Kar, K., Bashivan, P., Prescott-Roy, J., Schmidt, K., Yamins, D. L. K., & DiCarlo, J. J. (2018). Brain-Score: Which Artificial Neural Network for Object Recognition is most Brain-Like? (p. 407007). bioRxiv. 10.1101/407007

Serre, T. (2019). Deep Learning: The Good, the Bad, and the Ugly. Annual Review of Vision Science, 5(1), 399–426. 10.1146/annurev-vision-091718-014951

Shi, J., Shea-Brown, E., & Buice, M. A. (2019). Comparison Against Task Driven Artificial Neural Networks Reveals Functional Organization of Mouse Visual Cortex. *arXiv:1911.07986 [q-Bio]*. http://arxiv.org/abs/1911.07986

Spearman, C. (1904). The proof and measurement of association between two things. The American Journal of Psychology, 15(1), 72–101. 10.2307/1412159

Storrs, K. R., Kietzmann, T. C., Walther, A., Mehrer, J., & Kriegeskorte, N. (2021). Diverse Deep Neural Networks All Predict Human Inferior Temporal Cortex Well, After Training and Fitting. Journal of Cognitive Neuroscience, 1–21. 10.1162/jocn_a_01755

Teney, D., Nicolicioiu, A., Hartmann, V., & Abbasnejad, E. (2024). *Neural Redshift: Random Networks are not Random Functions* (arXiv:2403.02241). arXiv. 10.48550/arXiv.2403.02241

Tong, Z., & Tanaka, G. (2018). Reservoir Computing with Untrained Convolutional Neural Networks for Image Recognition. 2018 24th International Conference on Pattern Recognition (ICPR), 1289–1294. 10.1109/ICPR.2018.8545471

Wang, A. Y., Kay, K., Naselaris, T., Tarr, M. J., & Wehbe, L. (2023). Better models of human high-level visual cortex emerge from natural language supervision with a large and diverse dataset. Nature Machine Intelligence, 5(12), 1415–1426. 10.1038/s42256-023-00753-y

Yamins, D. L. K., & DiCarlo, J. J. (2016). Using goal-driven deep learning models to understand sensory cortex. Nature Neuroscience, 19(3), 356–365. 10.1038/nn.4244

Yamins, D. L. K., Hong, H., Cadieu, C. F., Solomon, E. A., Seibert, D., & DiCarlo, J. J. (2014). Performance-optimized hierarchical models predict neural responses in higher visual cortex. Proceedings of the National Academy of Sciences, 111(23), 8619–8624. 10.1073/pnas.1403112111

Yue, X., Pourladian, I. S., Tootell, R. B. H., & Ungerleider, L. G. (2014). Curvature-processing network in macaque visual cortex. PNAS Proceedings of the National Academy of Sciences of the United States of America, 111(33), e3467–e3475. 10.1073/pnas.1412616111

Yue, X., Robert, S., & Ungerleider, L. G. (2020). Curvature processing in human visual cortical areas. NeuroImage, 222. 10.1016/j.neuroimage.2020.117295

Zhou, B., Lapedriza, A., Khosla, A., Oliva, A., & Torralba, A. (2018). Places: A 10 Million Image Database for Scene Recognition. IEEE Transactions on Pattern Analysis and Machine Intelligence, 40(6), 1452–1464. IEEE Transactions on Pattern Analysis and Machine Intelligence. 10.1109/TPAMI.2017.2723009

